# Tiled Amplicon Sequencing Enables Culture-free Whole-Genome Sequencing of Pathogenic Bacteria From Clinical Specimens

**DOI:** 10.1101/2024.12.19.629550

**Authors:** Chaney C Kalinich, Freddy L Gonzalez, Alice Osmaston, Mallery I Breban, Isabel Distefano, Candy Leon, Patricia Sheen, Mirko Zimic, Jorge Coronel, Grace Tan, Valeriu Crudu, Nelly Ciobanu, Alexandru Codreanu, Walter Solano, Jimena Ráez, Orchid M Allicock, Chrispin Chaguza, Anne L Wyllie, Matthew Brandt, Daniel M Weinberger, Benjamin Sobkowiak, Ted Cohen, Louis Grandjean, Nathan D Grubaugh, Seth N Redmond

**Affiliations:** Department of Epidemiology of Microbial Diseases, Yale School of Public Health, New Haven, Connecticut, USA; Department of Ecology and Evolutionary Biology, Yale University, New Haven, Connecticut, USA; Department of Infection, Immunity, and Inflammation, Institute of Child Health, University College Longon, London, England; Universidad Peruana Cayetano Heredia, Lima, Peru; Institute of Phthisiopneumology, Chisinau, Moldova; Yale Institute for Global Health, Yale University, New Haven, Connecticut, USA; Public Health Modeling Unit, Yale School of Public Health, New Haven, Connecticut, USA

## Abstract

Pathogen sequencing is an important tool for disease surveillance and demonstrated its high value during the COVID-19 pandemic. Viral sequencing during the pandemic allowed us to track disease spread, quickly identify new variants, and guide the development of vaccines. Tiled amplicon sequencing, in which a panel of primers is used for multiplex amplification of fragments across an entire genome, was the cornerstone of SARS-CoV-2 sequencing. The speed, reliability, and cost-effectiveness of this method led to its implementation in academic and public health laboratories across the world and adaptation to a broad range of viral pathogens. However, similar methods are not available for larger bacterial genomes, for which whole-genome sequencing typically requires *in vitro* culture. This increases costs, error rates and turnaround times. The need to culture poses particular problems for medically important bacteria such as *Mycobacterium tuberculosis,* which are slow to grow and challenging to culture. As a proof of concept, we developed two novel whole-genome amplicon panels for *M. tuberculosis* and *Streptococcus pneumoniae*. Applying our amplicon panels to clinical samples, we show the ability to classify pathogen subgroups and to reliably identify markers of drug resistance without culturing. Development of this work in clinical settings has the potential to dramatically reduce the time of diagnosis of drug resistance for multiple drugs in parallel, enabling earlier intervention for high priority pathogens.

## Introduction

In recent years, whole-genome amplicon sequencing has been adopted as a standard technique for genomic surveillance of viral pathogens. Initially developed for genomic surveillance of the 2016 Zika epidemic [1], where low viraemia had precluded direct sequencing of clinical samples even where the infection had been confirmed, amplicon sequencing uses multiplex PCR of tiled overlapping regions of a target genome to recover whole genomes from samples of low concentration or complex microbial communities. This has proven particularly useful for sequencing remnant samples from diagnostic tests, and its use in the ‘ARTIC’ protocol for sequencing SARS-CoV-2 [2] has led to it being deployed in thousands of public health laboratories around the world, facilitating true global surveillance of viral dynamics [3]. The ease, reliability, and low cost of amplicon sequencing have seen its adaptation to a broad range of viral pathogens both in respiratory disease [4,5] and beyond [6,7]. However, more than half of all infectious disease-related deaths are caused by 33 bacterial pathogens [8]. Of these, *Mycobacterium tuberculosis* is the world’s most deadly single pathogen, responsible for 1.5 million deaths annually [9]. COVID-19-related disruptions are estimated to be responsible for approximately 700,000 excess tuberculosis (TB) deaths in the period 2020-2023, underscoring how vital it is to transfer advances in public health infrastructure related to the SARS-CoV-2 pandemic to make up for lost ground in controlling TB [9].

Whole genome sequencing (WGS) has proven invaluable for both viral and bacterial pathogens, enabling the reconstruction of transmission chains and routine genomic surveillance [10–13]. Similar to viruses, for bacteria such as *M. tuberculosis* and *Streptococcus pneumoniae*, WGS can facilitate targeting of interventions, including vaccines and screening programs [14–16]. In addition, WGS of bacteria offers unique advantages for detecting antimicrobial resistance and informing tailored treatment regimens [17–19]. WGS of *M. tuberculosis* in particular has also demonstrated clear application to detecting superspreading [20], or distinguishing recrudescence from reinfection [21].

Despite the demonstrated value of genomic surveillance, limitations of existing genomic tools have restricted their use to high-resource settings and/or retrospective analyses. *M. tuberculosis* has low genomic diversity, making traditional approaches, such as restriction fragment length polymorphism (RFLP), spoligotyping, and variable number tandem repeat (VNTR) insufficient to infer transmission in many circumstances [22]. Furthermore, these approaches do not provide information on drug resistance. Previous approaches to bacterial WGS rely on culture, which is relatively laborious and time-consuming. For *M. tuberculosis* in particular, a slow generation time of 24 hours leads to delays of up to 4-6 weeks before growth is observed on solid media [22]. Though sputum samples, having high viscosity and relatively low bacterial load, can be handled in many local health centers, the higher risk of airborne contagion in bacterial culture necessitates greater biosafety infrastructure and expertise, and is typically handled in a regional reference laboratory [23]. Assembling resources for culturing, extraction, and sequencing can be particularly challenging in the LMIC settings where TB is most prevalent. The use of WGS to inform outbreak investigations as they occur has therefore been limited to high-resource settings such as the United Kingdom [24]. Amplicon sequencing offers a potential solution. Typically used for viral pathogens, this approach allows the recovery of whole genomes from clinical samples, without the need for culturing. Adapting these techniques to bacterial pathogens would provide rapid culture-free sequencing from minimal input volumes. This could unlock the use of remnant tests, such as GeneXpert cartridges [25] for *M. tuberculosis*, or clinical swabs for undiagnosed respiratory tract infections, which are often collected in antibiotic-containing viral transport media but may contain pathogenic bacteria such as *S. pneumoniae*. The capacity to use such passive surveillance techniques could be transformative for bacterial genomic epidemiology.

We present here the first use of amplicon-based WGS for the sequencing of bacteria. We have selected two bacterial pathogens, *M. tuberculosis* and *S. pneumoniae* serotype 3, each of which impose a significant global burden of respiratory disease, yet differ markedly in epidemiology, transmission, evolutionary dynamics, and genome structure. Our tiled amplicon schemes are able to generate complete genome coverage from samples with minimal input concentrations without any requirement for bacterial culturing. We compare recovery of genomic data from a broad range of sample types, including saliva, sputum, and remnant clinical samples from diagnostic testing, and further show that this genomic data can reliably perform *in silico* lineage or serotype assignment to enable the surveillance of bacterial transmission dynamics. We show that our *M. tuberculosis* amplicon panel can be applied directly to sputum samples to identify clinically relevant phenotypes, such as antimicrobial susceptibility, within days of sample collection and can detect resistance loci that were not found by rapid diagnostics. We hope that this work will not only generate opportunities for future genomic epidemiology of *M. tuberculosis*, but will also provide a roadmap for the development of amplicon sequencing for other clinically important bacterial pathogens.

## Results

### *In silico* predictions indicate applicability of amplicon schemes to large bacterial genomes

In order to design primer schemes with efficient amplification of diverse target sequences, we downloaded a selection of whole-genome sequences available in public repositories for both *M. tuberculosis* (n=489, File S1A) and *S. pneumoniae* (n=490, File S1B). For *S. pneumoniae*, which is highly recombinogenic and frequently reshuffles its genome via horizontal gene transfer, we intended to target the primers in the core genomic regions shared across several strains rather than the ever-changing accessory genomic regions unique to individual strains. We achieved this by assembling these sequences into a reference-guided pseudo-whole-genome alignment based on genomes from multiple strains mapped to a serotype 3 reference and then generated consensus sequences or ‘metaconsensus’ to use for primer design. PrimalScheme [1] was run on these reference-guided aligned consensus sequences, in order to design primers covering the core genomic regions of *S. pneumoniae* genome as well as some partially conserved accessory genomic regions while avoiding highly variable sites. Because *M. tuberculosis* has very little within-species diversity outside of the repetitive PE/PPE regions (conserved proline-glutamine/proline-proline-glutamine domains and hypervariable C termini [26]), PrimalScheme was run directly on the H37Rv reference genome (NCBI accession NC_000962.3) after masking PE/PPE regions and sites with known resistance-related polymorphisms to avoid primers being designed at these loci. *S. pneumoniae*, which has an average core genome size of 2,160,000 bp [27], required 2292 primers to cover the entire core genome while *M. tuberculosis*, which has an average genome size of 4,411,000 [28], required 5128 primers to amplify the entire genome.

For both pathogens, we selected a small number of genetically diverse sequences from the larger set of publicly available sequences to predict coverage beyond the sequences used for primer panel design (see Methods: Off-target amplification prediction). As expected, predicted amplicon coverage was highest against the strains used as a reference for panel design (*Sp*:OXC141: 98.93%; *Mt*/H37Rv: 94.31%) - *M. tuberculosis* coverage is reduced due to the omission of PE/PPE regions from the design, which account for 8-10% of the genome [18]. However, predicted coverage in *M. tuberculosis* remained high across all members of the *M. tuberculosis* complex, including all 7 major lineages and *Mycobacterium bovis* (>= 94.23%), as well as *Mycobacterium canettii* (89.44%), while coverage of *S. pneumoniae* fell sharply between Global Pneumococcal Sequence Clusters (GPSCs) ( <82.00%->88.00%) and in the *Streptococcus mitis* outgroup (32.18%) due to the extremely high genetic diversity of the *S. mitis* lineages in the Mitis species complex (Figure 1) [29]. The range of average nucleotide identity was narrower across *M. tuberculosis* (99.98-99.27%) compared to the broader variation observed among *S. pneumoniae* (98.76-92.51%), reflecting the diversity in the GPSC lineages included. Pangenome size and similarity were also markedly different between the two species, with *M. tuberculosis* having a smaller relative pangenome (4,335 total genes and a mean genome size of 4,067 genes, a ratio of 1.07, based on the 13 genomes included) than *S. pneumoniae* (3,942 total genes and a mean genome size of 2,071 genes, a ratio of 1.90, based on the 13 genomes included). As a result, far more sharing of genes with the reference strain was observed for *M. tuberculosis* (4,002-4,050 shared genes) than *S. pneumoniae* (1,705-1,793 shared genes). This difference reflects the open nature of the *S. pneumoniae* pangenome, which facilitates adaptive responses by accommodating genetic diversity in a range of environments [30]. Our findings suggest that panel applicability is influenced by both genome rearrangements and increases in genetic distance, as evidenced by the low predicted coverage observed in cases of high genetic distance, such as between *S. pneumoniae* and *S. mitis*.

**Figure 1:**
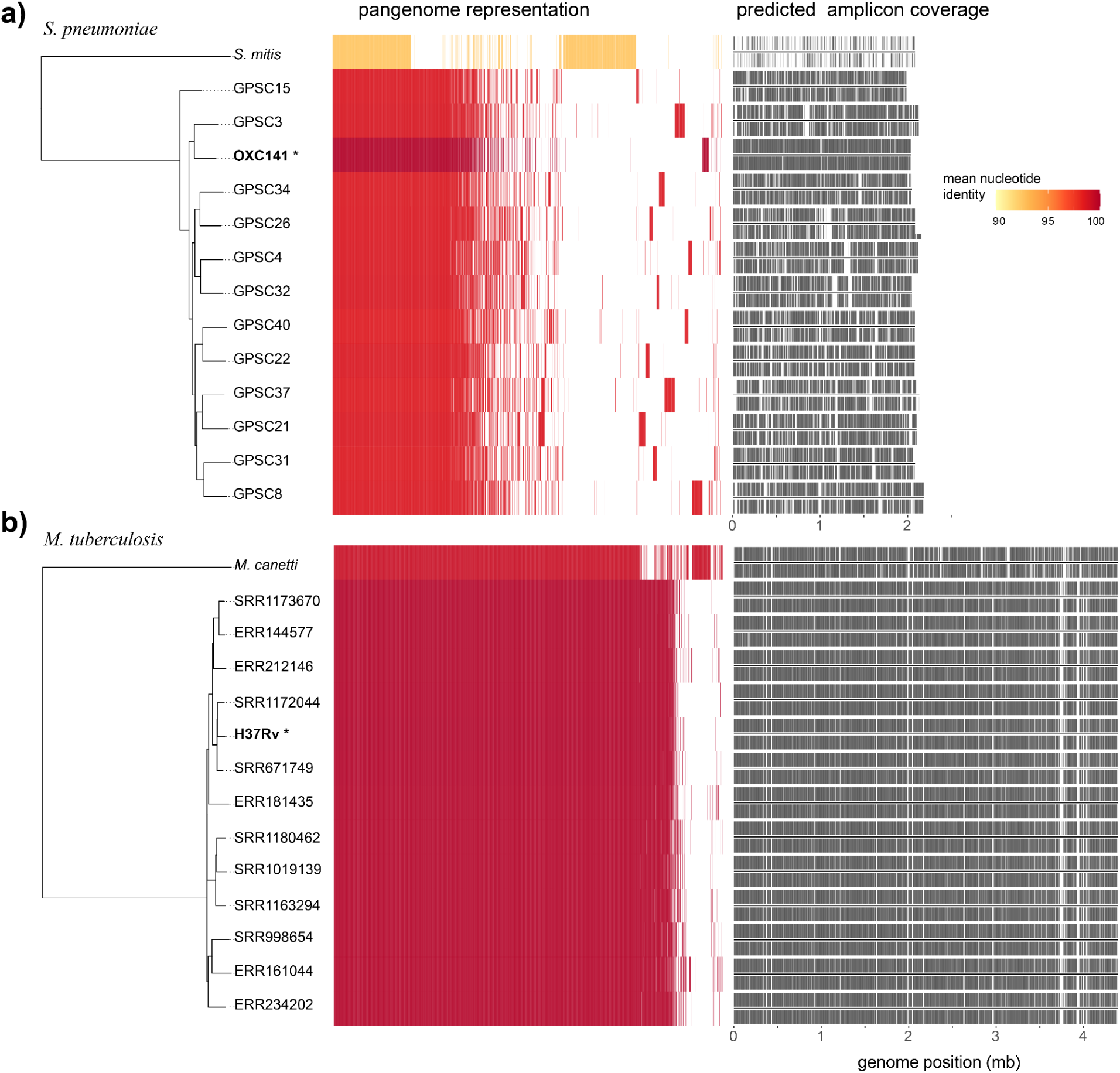
*In silico* modeling indicates broad applicability across diverse *Streptococcus pneumoniae* serotypes and *Mycobacterium tuberculosis* lineages. Pangenome representation of (A) *S. pneumoniae* whole genome sequences (n=13) and *Streptococcus mitis* outgroup (Accession: AP023349) and (B) *M. tuberculosis* whole genome sequences (n=13) and *Mycobacterium canettii* outgroup (Accession: NC_019950). Starred phylogenetic tree tips mark the reference sequences used for primer design. Shaded bar graphs (middle) denote genes shared amongst clades, yellow-red color scale denotes average nucleotide identity. Predicted amplicon coverage (right) is shown in grey with forward and reverse amplicon pairs displayed above and below the line. *M. tuberculosis* has a higher proportion of genes shared between clades and higher ANI than the open pangenome of *S. pneumoniae*, driving the higher predicted amplicon coverage. A list of the sequences used in this analysis can be found in Table S2.

### Amplicon sequencing enables recovery of *M. tuberculosis* whole genomes from clinical specimens without prior culturing

For *M. tuberculosis*, we sequenced DNA from cultured isolates and sputum samples using a standard amplicon sequencing workflow, with and without adding amplicon panel primers. Comparisons of amplified and unamplified *M. tuberculosis*-positive sputum samples demonstrated dramatic increases in coverage for amplified samples as compared to the same samples without amplification (Figure 2A-B). While only 2/10 of unamplified samples achieved more than 75% coverage, 9/10 of the amplified samples sequenced above this threshold, with 7 of those generating more than 95% coverage. The remaining sample achieved 33% coverage amplified, and negligible coverage unamplified. Metagenomic sequencing indicated successful amplification from samples containing both commensal and pathogenic bacteria including *Streptococcus, Pseudomonas, Actinomycetes* and *Schaalia spp*.

**Figure 2:**
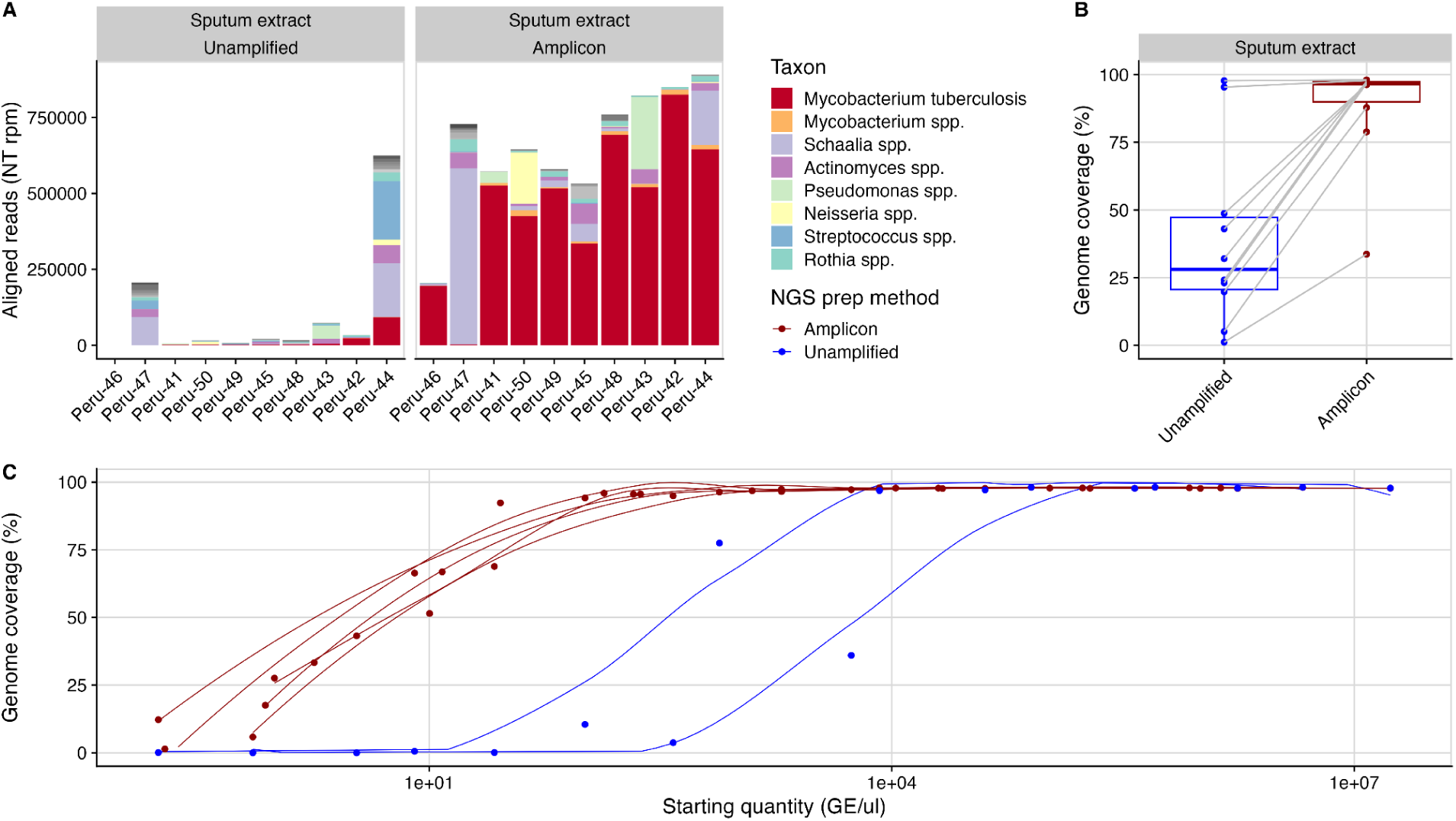
Tiled amplicon sequencing enables culture-free recovery of whole genome sequences from *Mycobacterium tuberculosis* sputum specimens. We compared sequencing results for unamplified and amplified *M. tuberculosis* clinical specimens with regard to (A) microbial content via the CZID metagenomics pipeline and (B) average genome coverage. *M. tuberculosis*-positive sputum samples sequenced directly showed dramatic increases in genome coverage, with 8/10 samples generating more than 80% coverage after amplification with our protocol, and a ninth sample generating 78% coverage despite a significant infection with *Schaalia odontolytica*. Serial dilutions of DNA from cultured isolates (C) demonstrated that the amplicon scheme enables recovery of genomes with >95% coverage from all samples with ≥100 GE/μL, compared to ≥10,000 GE/μL without amplification.

We assessed the limits of detection for each amplicon panel by sequencing serial 10-fold dilutions of 6 cultured isolates using both amplified and unamplified sequencing approaches. For *M. tuberculosis*, high genome coverage (>95%) was observed in all amplified samples above 100 genome equivalents per microliter (GE/µL), compared to 10,000 GE/µL for unamplified samples (Figure 2, S5).

### Direct sputum sequencing enables phylogenetic classification and detects markers of antimicrobial resistance for *M. tuberculosis*

Lineages of *M. tuberculosis* were called with the Mykrobe package [31], which assigned all samples to lineages 2 (sublineage 2.2.1) or 4. Mykrobe performed equally well in high coverage samples, regardless of whether these were derived from cell culture or sputum. We derived maximum-likelihood (ML) phylogenies using IQ-tree [32] including all sequenced specimens and the broad reference set of *M. tuberculosis* sequences used for primer design (Figure S2; File S1A). In all cases the primary lineage predicted by Mykrobe aligned with lineages from an ML tree, though in some cases secondary lineages were predicted based on minor variants which did not concord with the ML tree.

For *M. tuberculosis*, sequencing data was high-quality enough to produce a prediction for all template dilutions from cultured isolates with at least 10 GE/µL starting quantity (Figure 4, Figure S3). While we do not have access to phenotypic susceptibility results for these isolates, genotypic-based predictions of drug resistance in *M. tuberculosis* are usually highly accurate for most drug classes [33]. These predictions were internally consistent for all template dilutions above 100 GE/µL (though there was some variability between partial vs. full resistance calling) with the exception of streptomycin. DNA was extracted directly, without culturing, from 60 sputum specimens with a range of acid-fast bacilli semi-quantitative measurements (e.g., 1+ to 3+). Sequencing data was high-quality enough to produce a drug susceptibility prediction for 53/60 sputum specimens. Of the seven specimens which failed, four (Yale-TB121, Yale-TB123, Yale-TB149, Yale-TB150) had starting quantities (following extraction) below 10 GE/µL. None of the other three (Yale-TB126, Yale-TB139, Yale-TB148) had GeneXpert results available as comparison. Several different extraction methods were used (detailed in Table S1B and Appendix S1) as it was not clear what method would perform best; all 10 specimens extracted with the final protocol, which included a NALC-NaOH treatment to deplete non-mycobacterial DNA, had adequate data to predict resistance.

### Tiled amplicon sequencing amplifies *S. pneumoniae* amongst mixed species clinical samples

To determine our ability to enrich target genomes from within diverse sample sources, for *S. pneumoniae*, we sequenced DNA from cultured isolates, saliva samples, and culture-enriched saliva, both using our amplicon panel and standard metagenomic sequencing. Despite expected high complements of *Streptococcus oralis* and *S. mitis* in saliva samples, post-amplification samples showed an increased proportion of reads mapping to the *S. pneumoniae* target genome compared to standard metagenomic sequencing. However, this was accompanied by an increase in reads mapping to related *Streptococcus* species indicating some non-specific amplification across the *Streptococcus* genus (Figure 3A-B). Serial dilutions of cultured isolates indicate a low limit of detection with recovery of > 95% of the genome in most (7/8) samples above 1,000 GE/μL, a 50-fold increase over unamplified samples which were universally unsequenceable below 50,000 GE/μL (Figure 3C).

**Figure 3.**
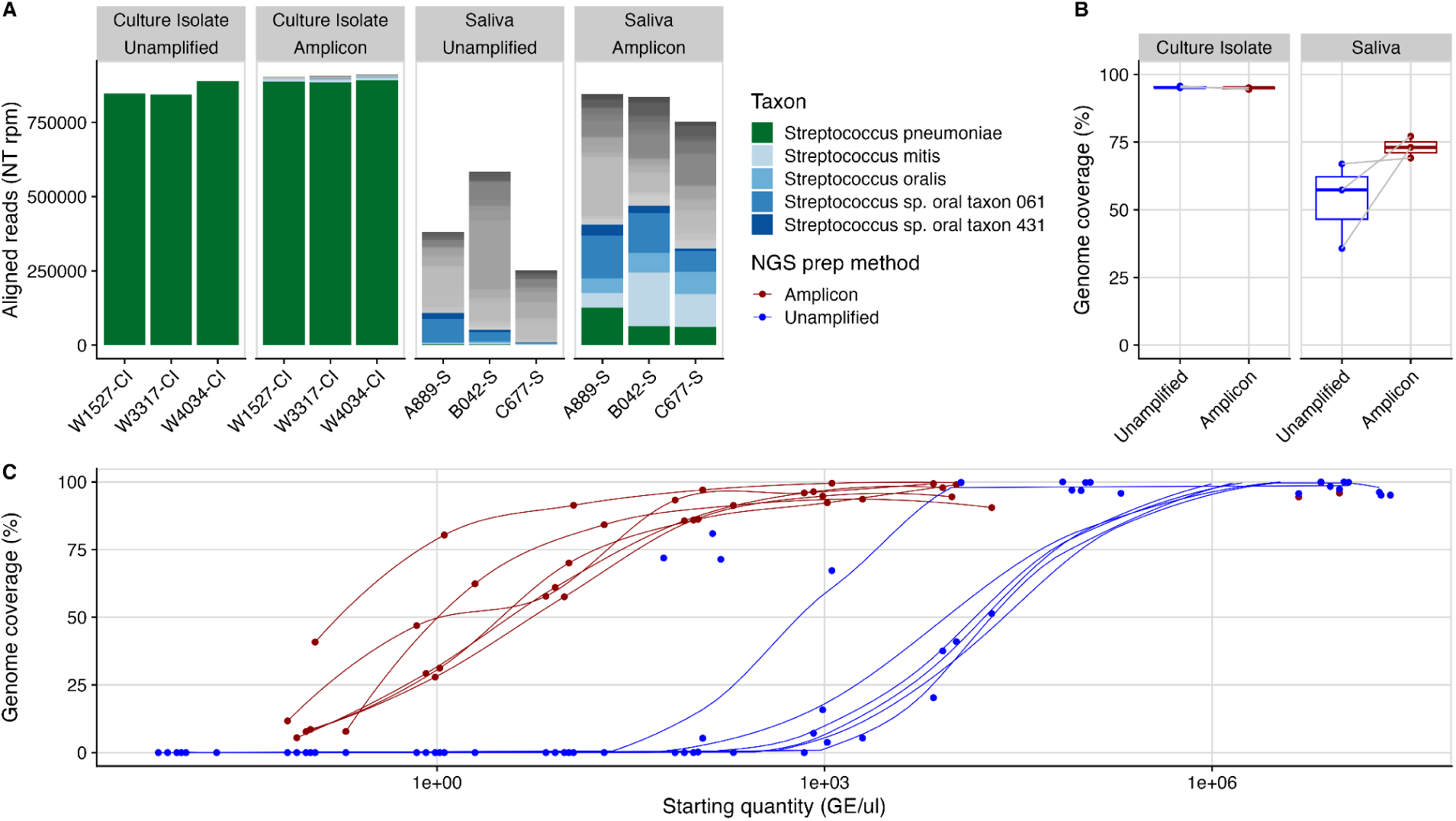
Tiled amplicon sequencing enhances culture-free recovery of whole genome sequences from S. pneumoniae sputum specimens. We compared sequencing results for amplified and unamplified *S. pneumoniae* clinical specimens with regard to (A) microbial content via the CZID metagenomics pipeline and (B) average genome coverage. *S. pneumoniae*-positive respiratory samples from matched patients showed increases in genome coverage and depth for all sample types, despite simultaneous amplification of closely related taxa. Non-Streptococcus taxa are indicated by gray shading. Serial dilutions of DNA from cultured isolates (C) demonstrated that the amplicon scheme enables recovery of genomes with >95% coverage from most (7/8) isolates with starting quantities ≥1,000 GE/μL, while no isolates below 10,000 GE/μL achieved this without amplification.

**Figure 4:**
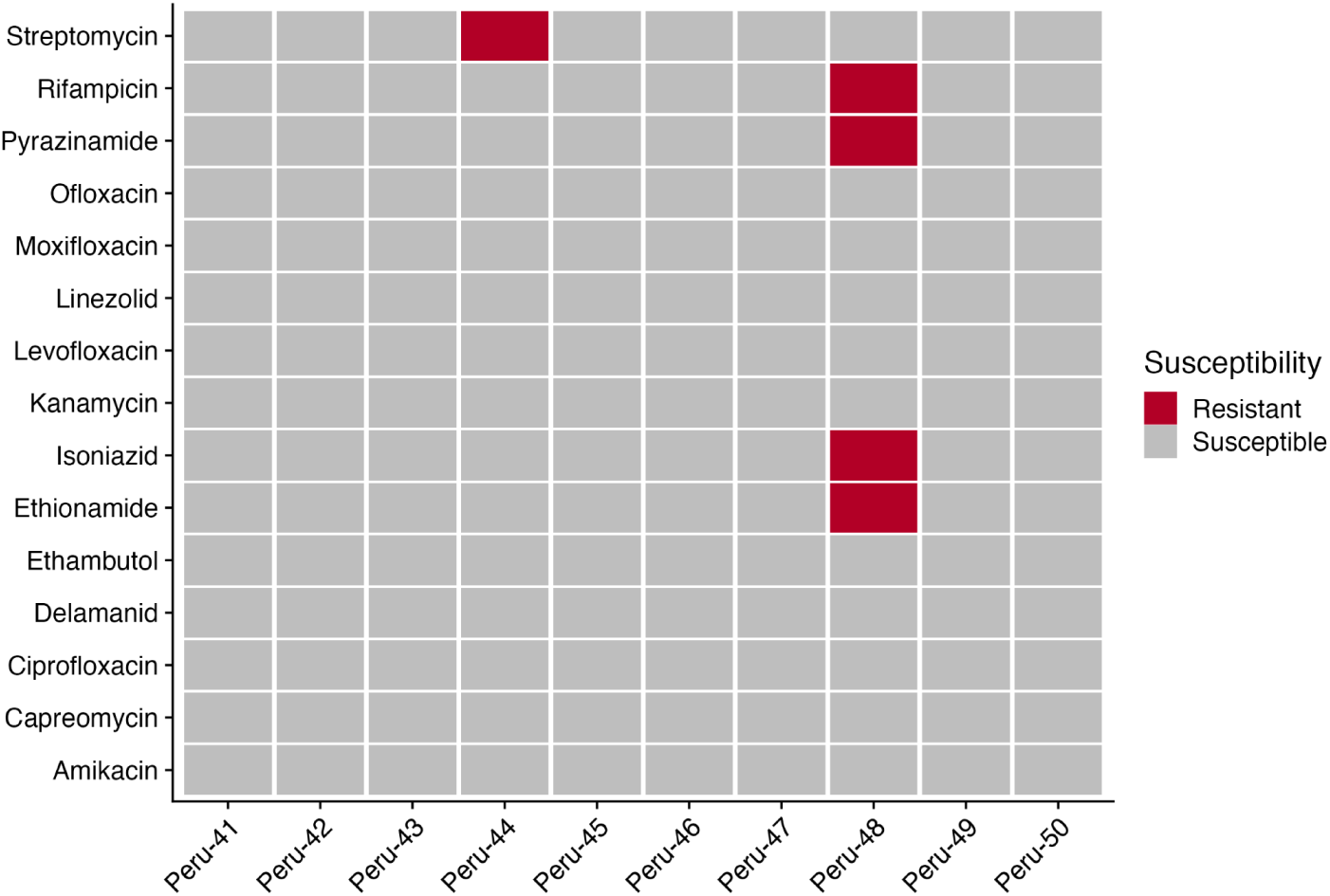
Amplicon sequencing predicts TB antimicrobial resistance *in silico*. Predicted susceptibility to 15 anti-TB drugs by amplicon sequencing for DNA extracted from sputum without prior culture using our optimized extraction protocol, showing detection of Streptomycin and combined Rifampicin / Isoniazid resistance.

### Phylogenetic classification and antimicrobial resistance is not readily assessed using amplicon derived data from *S. pneumoniae*

Lineage calling for *S. pneumoniae* using the GPSC nomenclature [34], implemented in the PopPUNK framework [35], was largely unsuccessful. Culture-derived samples could be assigned to capsular serotype using PneumoKITy [36] (Table S3), however none of the lineage callers (PopPunk [35] and SRST2 [37]) were able to assign lineages to any of the direct clinical samples from saliva or culture isolates. While our data did indicate coverage of the 7 major *S. pneumoniae* housekeeping genes (*aroE, gdh, gki, recP, spi, xpt, ddl*) used to assign sequence types (STs) or clones based on pneumococcal multilocus sequence typing (MLST) scheme [38] (Table S4; Figure S4), lineage or clone predictions may have been impaired by the high concentration of commensal bacteria in the enriched samples which harbor similar genes in their chromosomes. Further development is required to design primers which are specific to pathogenic *Streptococcus* strains.

*In silico* analysis identified antibiotic resistance against several second-line and broad-spectrum antibiotics, including Lincosamides, Macrolides, and Fluoroquinolones (Table S5). Resistance was detected in 8/9 culture isolate samples, 2/3 raw saliva samples, and 1/3 culture enriched saliva samples. Percent coverage and identity for resistance genes (*RlmA(II), patA, patB, and pmrA*) ranged from 88.33% to 100% (mean = 99.57%) and 99.25% to 100% (mean = 99.58%) for cultured isolates, and from 78.72% to 100% ( mean = 94.28%) and 84.32% and 94.36% (mean = 89.90%) in raw saliva. The single culture enriched saliva sample showed 87.75% coverage and 85.77% percent identity toward *RlmA(II).* It is worth noting that the presence of antibiotic resistance genes in samples containing mixed *Streptococcus* species introduces potential challenges in definitely attributing resistance determinants to *S. pneumoniae* without additional confirmatory analyses.

## Discussion

Tiled amplicon sequencing of pathogens has proven extremely useful for reconstructing disease spread and gaining insight into transmission patterns for a variety of viruses [39]. The COVID-19 pandemic stimulated a global effort to adopt these methods and use genomics to track and monitor SARS-CoV-2; however, it has not previously been applied to the significantly larger and often more complex genomes of bacteria. We report the first use of a tiled amplicon approach to sequence two pathogenic bacteria from specimens with minimal input DNA and demonstrated the ability to identify lineages and markers of drug resistance, which could have a major impact on disease control.

Both *S. pneumoniae* and *M. tuberculosis* are pathogens of prime public health importance. *S. pneumoniae* is responsible for more than 800,000 deaths per year, with the majority of these resulting from respiratory tract infections in children under five [8]. With the exception of three years during the COVID-19 pandemic (2020-2022), TB has been the leading cause of death from a single infectious agent, causing more than a million deaths per year [40]. Despite the availability of vaccines, treatment, and significant funding [41], we continue to miss WHO targets for reductions in TB incidence and death by wide margins. Rather, cases have risen worldwide over the past 2 years [42]. Much of this rise is attributable to interruptions in disease detection and treatment during the COVID-19 pandemic. As of 2023, with *M. tuberculosis* reclaiming the title of most deadly single pathogen, there is a renewed focus on improving TB treatment and control - in particular, there are 6 vaccine candidates currently in Phase III clinical trials [43] and efforts to develop shorter and more effective treatment regimens [44]. This may be an ideal time to apply tools used to control and monitor interventions for COVID-19 to TB [9].

Antimicrobial resistance is a critical issue in treating and controlling TB, due to the prevalence of resistance to first line drugs and the length, cost, and complexity of treatment regimens [45]. Despite the introduction of shorter regimens, the time taken to find an effective treatment can be long, and incomplete treatment remains a problem [46]. For this reason, the WHO now recommends the use of targeted sequence-based diagnostics for rapid drug susceptibility testing for patients who are at high risk of, or have already experienced, treatment failure [47]. However, designing such an assay is not simple; more than 40 separate loci, each containing numerous individual mutations, have been implicated in drug resistance [48], and uncertainty can be higher for new or second line drugs [49]. WGS works around these limitations of targeted amplicon sequencing. Unlike targeted sequencing approaches, data are being generated across the entire genome, therefore as new genetic markers of resistance are discovered, these can be added bioinformatically without adding new primers. The required time, infrastructure, and costs for tiled amplicon sequencing are almost identical to targeted amplicons; the additional data generated through WGS can be used along with phenotypic drug susceptibility to expand our understanding of the genetic markers of drug resistance, especially for third-line or novel drugs, increasing the accuracy of predictions over time [18].

WGS obviates the need to design a targeted assay and can also return resistance predictions within days of a positive culture. However, the requirement of most existing WGS approaches to first culture a sample means that the overall sample-to-sequence turnaround time for *M. tuberculosis* is measured in weeks or months [24] and significant biases can be introduced during the culturing process itself [18]. Direct WGS without culture does not consistently produce data of high enough quality for resistance prediction or thorough epidemiologic investigation [50,51], is limited to specimens with a high bacterial load [51], or relies on expensive techniques such as hybrid capture [52]. We have demonstrated tiled amplicon sequencing directly from sputum specimens, without culture, can be used to make accurate drug susceptibility predictions and lineage assignments for the majority (53/60) of specimens, unlike prior whole-genome sequencing approaches [24,50–53]. For a notoriously slow-growing organism such as *M. tuberculosis*, eliminating this step reduces the time from sample collection to genome from weeks to days. Not only could patients receive appropriate antibiotics sooner, but genomic epidemiology could be used in real-time to inform outbreak investigations [10,54] and public health measures to reduce spread [11,55].

Neither panel showed high rates of amplicon dropout when faced with targets which had drifted from the reference strain (a regular issue with viral amplicon panels). However, *in silico* predictions suggest a weaker applicability of the amplicon panel in species which undergo significant levels of recombination and horizontal gene transfer, and our inability to reliably recover serotype and resistance loci in *S. pneumoniae* supports this conclusion. Indeed, *S. pneumoniae* exhibits extremely high levels of horizontal gene transfer, not only within the species, but also with frequently co-occurring commensal bacteria such as *S. mitis* and *S. oralis* [56,57]. This makes it a particularly challenging target for whole-genome reconstruction using amplicon-based approaches. If the goal is to determine *S. pneumoniae* serotype, antimicrobial resistance, or lineage, an amplicon panel specifically targeted to the relevant genomic regions may be more reliable. *M. tuberculosis* showed consistently high coverage across both *in vitro* and *in silico* predictions, however gaps remain in our coverage of the genome in the PE/PPE regions. While these are frequently omitted from *M. tuberculosis* analyses, increasing evidence of functions in host cell invasion [58] and importance for vaccine design [59] suggest inclusion of these regions in future iterations of this amplicon panel would be a significant improvement.

The comparison between our results for *S. pneumoniae* and *M. tuberculosis* is instructive. Faced with both drift and genomic rearrangement, designing primers that target conserved motifs will rely upon databases of previously sequenced genomes to allow us to determine circulating genetic diversity. Rapid improvements in sequencing and assembly technology have generated vast databases of assembled genomes; while these resources are not comprehensive, their bias towards improved representation of species of clinical interest [60] suggests this will not be a limiting factor in panel design. An alternative consideration is targeting bacteria which do not undergo significant levels of horizontal gene transfer, and the ratio of genome to pangenome size is likely to be a key metric for our ability to design an amplicon panel. This ratio is highly sensitive to the diversity of habitats in which the pathogen is found: free-living or commensal species, such as *S. pneumoniae*, gain particularly large pangenomes to enable adaptation to diverse environments; intracellular pathogens show strong purifying selection, low effective population sizes and low genome-to-pangenome ratios [61]. *M. tuberculosis*, an obligate pathogen and intracellular bacterium which has been extensively sequenced [19], has little horizontal gene transfer, and remains a major threat to human life [42], may be archetypal, however other intracellular pathogens such as *Yersinia pestis*, *Listeria monocytogenes*, *Legionella pneumophila*, *Chlamydia trachomatis, Treponema pallidum, and Neisseria Gonorrhoeae* are suitable targets.

The widespread use of tiled amplicon sequencing for pathogen genomics during the COVID-19 pandemic has ensured that this method is trusted, understood, and easily implemented in academic and public health laboratories worldwide. As the focus now turns to adapting this capacity to other public health threats [3], it is important to prioritize the development of tools for global priority pathogens that can be implemented in the regions suffering the greatest burden. The method presented here uses the exact same workflow as used by thousands of public health labs to sequence SARS-CoV-2, utilizing an off-the-shelf commercial sequencing library preparation kit with the single change of swapping out primer pools used to generate amplicons. Genomic surveillance of *M. tuberculosis* has demonstrated capacity to guide TB interventions in high income countries [20,21]; the reductions in cost and turnaround time afforded by tiled amplicon sequencing could enable this to be implemented in LMICs with high TB burden. Just four countries (India, Bangladesh, Indonesia, Democratic Republic of the Congo) account for over half of all TB deaths; all four have prior in-country experience with amplicon sequencing of SARS-CoV-2 [62–65] suggesting a ready capacity for tiled amplicon sequencing of *M. tuberculosis*. Extensive use of alternative sequencing methods such as Oxford Nanopore in these regions [62,64,66] suggest adaptation to cheaper and more portable sequencing platforms may further increase surveillance capacity.

Barriers to clinical application are necessarily higher [67]. If diagnostics and resistance prediction are to be used to tailor treatment regimens, it is vital that they can be shown to work reliably in a range of likely scenarios: paucibacillary infections; mixed *M. tuberculosis* strains; mixed *M. tuberculosis* and nontuberculous mycobacteria; partial and incomplete resistance. A comprehensive evaluation of culture-free sequencing methods in a clinical environment should be a priority for TB control, permitting earlier diagnosis and resistance profiling. Despite this complex landscape, the capacity of culture-free *M. tuberculosis* sequencing could be transformative - not only by increasing rates of successful TB treatment at the patient level [68], but by preventing the further transmission of MDR-TB at the population level [69,70]. With several new TB vaccines in Phase III trials, WGS can also provide lineage-specific estimates of vaccine efficacy and early signal of vaccine escape. Optimizing the use of available antimicrobials and vaccines, both old and new, is vital to improve TB control and culture-free WGS can be used to ensure the right interventions are reaching the right populations in a timely manner.

## Materials and Methods

### Ethics statement

All specimens were de-identified, remnant specimens used previously for diagnostic testing or IRB-approved human subjects research in accordance with Yale University IRB-exempt protocol #2000033281. *S. pneumoniae*-positive specimens were remnant specimens collected from study participants enrolled and sampled in accordance with the Yale University Humans Investigation Committee-approved protocol #2000027690. *M. tuberculosis*-positive specimens from Moldova were remnant specimens collected from study participants enrolled and sampled in accordance protocol #2000023071 approved by Yale University Human Investigations Committee and the Ethics Committee of Research of the Phthisiopneumology Institute, Moldova. *M. tuberculosis* specimens from Peru were remnant specimens collected from study participants enrolled in accordance with protocol #204749 approved by the Institutional Committee on Research Ethics at Cayetano Heredia University, Peru.

### Primer design

We downloaded all available *S. pneumoniae* serotype 3 contigs (n=490; File S1B) from the Global Pneumococcal Sequencing (GPS) database [34], filtered specifically for phenotypic serotype 3, as of February 2, 2023. We downloaded raw reads for *M. tuberculosis* sequences from a previously described globally representative dataset [19] (n=489; File S1A) from the European Nucleotide Archive at the European Molecular Biology Laboratory-European Bioinformatics Institute (EMBL-EBI). For both targets, we downloaded complete reference genomes from the National Center for Biotechnology Information (NCBI) GenBank (OXC141; accession NC_017592 and H37Rv; accession NC_000962.3).

For *M. tuberculosis*, variants were called against the reference using Snippy and a time-resolved ML tree was built using our variant call file, along with sample data generated from Augur (v.22.4.0) [71], IQ-Tree (v.2.23) [72], and TreeTime (v.0.10.1) [73]. Representative sequences (n=6) were selected from across this tree using Parnas (v.0.1.4) [74], to cover >50% of the expected overall diversity. We used these representatives to create an *M. tuberculosis* core genome assembly using Snippy.

For *S. pneumoniae*, consensus genome sequences were generated (n=4) with Snippy (v.4.6.0) (https://github.com/tseemann/snippy).

Tiled primer schemes (target amplicon size 2kb) were designed for both *S. pneumoniae* and *M. tuberculosis* (excluding PE/PPE and repeat regions) using PrimalScheme [1]. Primers were ordered at 100 µM and 200 µM in IDTE for *S. pneumoniae* and *M. tuberculosis*, respectively. Primer pools consisted of an equal volume of each primer and were used for amplification without further dilution.

### Clinical specimens

*S. pneumoniae* samples consisted of DNA extracted from raw saliva (15), culture-enriched saliva samples (16), and cultured pure isolates (9). All saliva specimens had a paired cultured specimen cultured from the saliva (either culture-enriched sample or cultured isolate). A full list of *S. pneumoniae* samples and descriptions can be found in Table S1A. DNA was extracted from 200 µL of each sample using the MagMAX Ultra viral/pathogen nucleic acid isolation kit (Thermo Fisher Scientific) using a KingFisher Apex instrument (Thermo Fisher Scientific) and quantified using two qPCR primer/probe pairs, *lytA* [75] and *piaB* [76], as described previously [77] (Table S6).

*M. tuberculosis* samples consisted of DNA extracted from positive solid or liquid cultures from sputum and DNA extracted directly from sputum specimens. Extracts from culture consisted of remnant specimens from a prior study in Moldova, where sputum specimens were tested at a number of diagnostic centers in Moldova by microscopy, Xpert, and culture and positive cultures sent to the National TB Reference Laboratory in Chisnau for extraction by the cetyltrimethylammonium bromide (CTAB) method, as described previously [11]. Extracts from sputum consisted of specimens collected in Peru after routine diagnostics had been carried out and the presence of *M. tuberculosis* confirmed. In order to test the efficiency of different methods for extracting DNA from sputum, each specimen was split into two and processed with two different protocols. A total of 30 unique sputum specimens were processed with two protocols each, and a total of 6 different protocols were tested. A full list of all *M. tuberculosis* samples and the extraction methods used can be found in Table S1B, and a detailed description of extraction methods can be found in Appendix S1. Following extraction, DNA was quantified with a *M. tuberculosis*-complex specific, fluorescence-based real-time PCR assay on the Bio-Rad CFX96 instrument [78].

### Metagenomic sequencing

For *S. pneumoniae*, 1-3 ng of each sample (up to 4 µL for samples which were undetectable) and a negative template control (4 µL nuclease-free water) underwent tagmentation for 5 minutes followed by a magnetic bead cleanup. Then, samples were amplified with Nextera dual-index adapters followed by a second magnetic bead cleanup. Each sample was quantified with a Qubit fluorometer and 5 ng of each library were pooled together (up to 4 µL for undetectable samples). The pooled libraries underwent a final 0.7X bead clean up, then were quantified on a Qubit fluorimeter and quality and fragment distribution verified using an Agilent Bioanalyzer. For *M. tuberculosis*, samples were prepared as described for amplicon sequencing, but with the addition of sterile water in place of PCR primer pools for both amplification reactions.

### Amplicon sequencing

Amplicon DNA was prepared using the Illumina COVIDSeq DNA prep kit with primer pools for either *S. pneumoniae* or *M. tuberculosis* alongside a negative template control, as previously described [79].

Template DNA from each specimen was amplified in two separate PCRs, one reaction for each primer pool. For each sample, equal amounts of each PCR product were combined and the 2 kb amplification products underwent tagmentation for 3 minutes followed by a bead cleanup and library amplification with Illumina index adapters. Equal volumes of the fragmented and indexed library for each sample was pooled, followed by size-selective bead cleanup for DNA fragments between 300-600 bp. The final pooled library was quantified with a Qubit fluorometer and dsDNA High-Sensitivity Assay kit, and the fragment distribution verified on an Agilent Bioanalyzer and high-sensitivity DNA kit. Pooled libraries were sequenced on an Illumina NovaSeq (paired-end 150) with an average of 10 million reads per library.

### Alignments & Calling

Reads were aligned to the appropriate reference (*S. pneumoniae*: CC180 (Serotype 3); *M. tuberculosis:* H37Rv) using BWA-MEM (v.2.2.1) [80] and SAMtools (v1.15.1) [81]. Amplicon sequencing data were filtered (using defaults; Q>20 over a sliding window of 4, minimum read length 50% of the average length). *M. tuberculosis* primer sequences were trimmed using iVar (v.1.4.2) [82]. Commonly used amplicon primer trimming software (iVar) showed little success at removing *S. pneumoniae* primers from the sequencing data, likely due to misalignment of primers with the target sequences. Metagenomic sequences were trimmed and filtered for quality and length (<100bp), using Trim Galore (v.0.6.10) [83]. Variants were called and filtered (Phred score Q>10 and read depth >10) using BCFtools (v.1.21) [84].

Read subsampling, depth, and coverage was calculated using SAMtools [81]. Raw reads were directly submitted to the CZID mNGS Illumina pipeline [85] for microbial composition characterization within samples. Further data analyses and visualizations were carried out in Rstudio (v.2024.04.2+764) [86] using the tidyverse suite (v.2.0.0) [87].

### Off-target amplification prediction

For each amplicon panel, off-target amplification was assessed *in silico* against a set of related genomes. We first downloaded *S. pneumoniae* complete genome assemblies (n=35) (Gladstone et al. 2019), representing 62% of the GPS database lineages. Lineages containing serotype 3 (i.e. GPSC 12) were excluded to demonstrate off-target amplification, though the reference genome for serotype 3 (i.e. OXC 141) was retained (Figure S1A). We similarly selected a subset of *M. tuberculosis* sequences (n=76) from our initial pool used to develop the *M. tuberculosis* primer set (Figure S1B). For each species, we further compiled a genome cluster consisting of the reference genome used for primer design, 12 diverse strains representing various lineages, and an outgroup (Table S2). The pangenome for each cluster was calculated using Roary (v.3.13.0) [88] and an ML phylogeny was constructed using FastTree (v.2.1.11) [89]. Average nucleotide identity was calculated between our references and all other genomes in the cluster using FastANI (v.1.34) [90]. Off-target amplification was inferred by primer alignment using Bowtie (v.1.3.1) [91]; amplicons were predicted for any properly oriented amplicon pairs within 2,200 bp.

### Serotyping, lineage assignment, and resistance prediction

*S. pneumoniae* raw reads were first filtered against OXC141 and later *de novo* assembled with the CZID mNGS Illumina pipeline [92]. *In silico* multi-locus sequencing types (MLST) were assigned with mlst (v.2.23.0) [93]. GPSCs were assigned with PopPUNK (v.2.7.0) [35]. *In silico* screening of contigs for *S. pneumoniae* antimicrobial and virulence genes was done using ABRicate (v1.0.1) [94] and appropriate antimicrobial resistance databases [95–98]. For *M. tuberculosis*, Mykrobe [31] was used to both assign lineages and predict resistance using the built-in panel “202309” for *M. tuberculosis* [99]. As a comparison, a time-resolved ML tree was built using our variant call file, along with sample data generated from Augur (v.22.4.0), IQ-Tree (v.2.23), and TreeTime (v.0.10.1) (Figure S2). Tree visualizations were done using Auspice (v2.57.0).

## Supporting information

Main supplement

