## Supplementary material for "Tiled Amplicon Sequencing Enables Culture-free Whole-Genome Sequencing of Pathogenic Bacteria From Clinical Specimens": Main supplement

### Contents

Figure S1A. Streptococcus pneumoniae pangenome. 5
Figure S1B. Mycobacterium tuberculosis pangenome. 5
Figure S2. Amplicon sequencing can be used in phylogenetic investigation. 6
Figure S3. Efficiency of extraction protocol affects antimicrobial resistance predictions. 7
Figure S4. Raw read alignments to Streptococcus pneumoniae serotype 3 housekeeping genes under amplified and unamplified sequencing conditions. 9
Table S1A. S. pneumoniae clinical specimens 10
Table S1B. M. tuberculosis clinical specimens. 11
Table S2. Samples used to predict off-target amplification. 15
Table S3. PneumoKITy analysis of S. pneumoniae. 16
Table S4. MLST analysis of S. pneumoniae culture isolates. 20
Table S5. Antibiotic resistance in S. pneumoniae samples. 21
Appendix S1. M. tuberculosis extraction methods from sputum 24

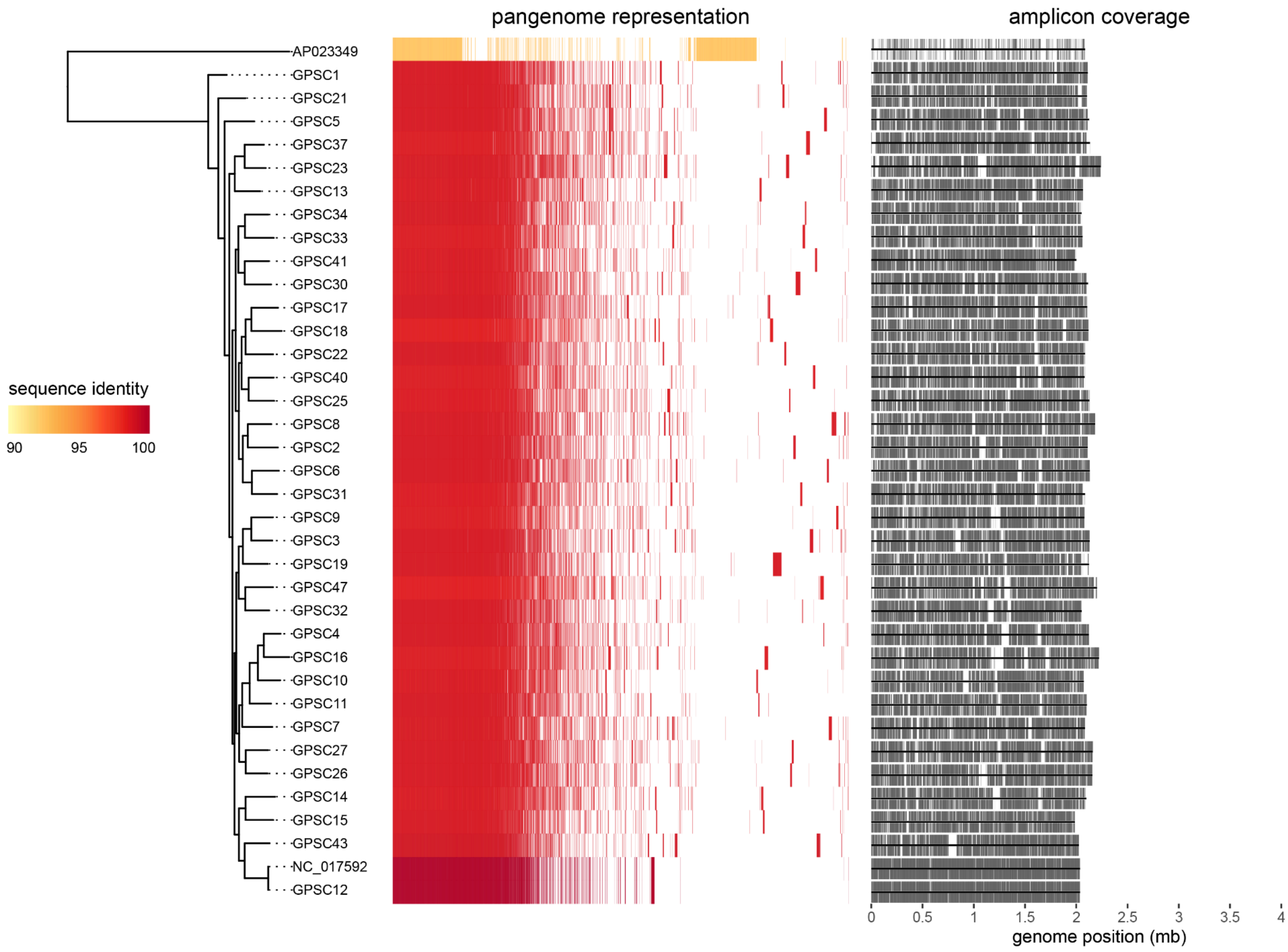

pangenome representation
amplicon coverage
sequence identity
90
95
100
AP023349
GPSC1
GPSC21
GPSC5
GPSC37
GPSC23
GPSC13
GPSC34
GPSC33
GPSC41
GPSC30
GPSC17
GPSC18
GPSC22
GPSC40
GPSC25
GPSC8
GPSC2
GPSC6
GPSC31
GPSC9
GPSC3
GPSC19
GPSC47
GPSC32
GPSC4
GPSC16
GPSC10
GPSC11
GPSC7
GPSC27
GPSC26
GPSC14
GPSC15
GPSC43
NC_017592
GPSC12
0
0.5
1
1.5
2
2.5
3
3.5
4
genome position (mb)

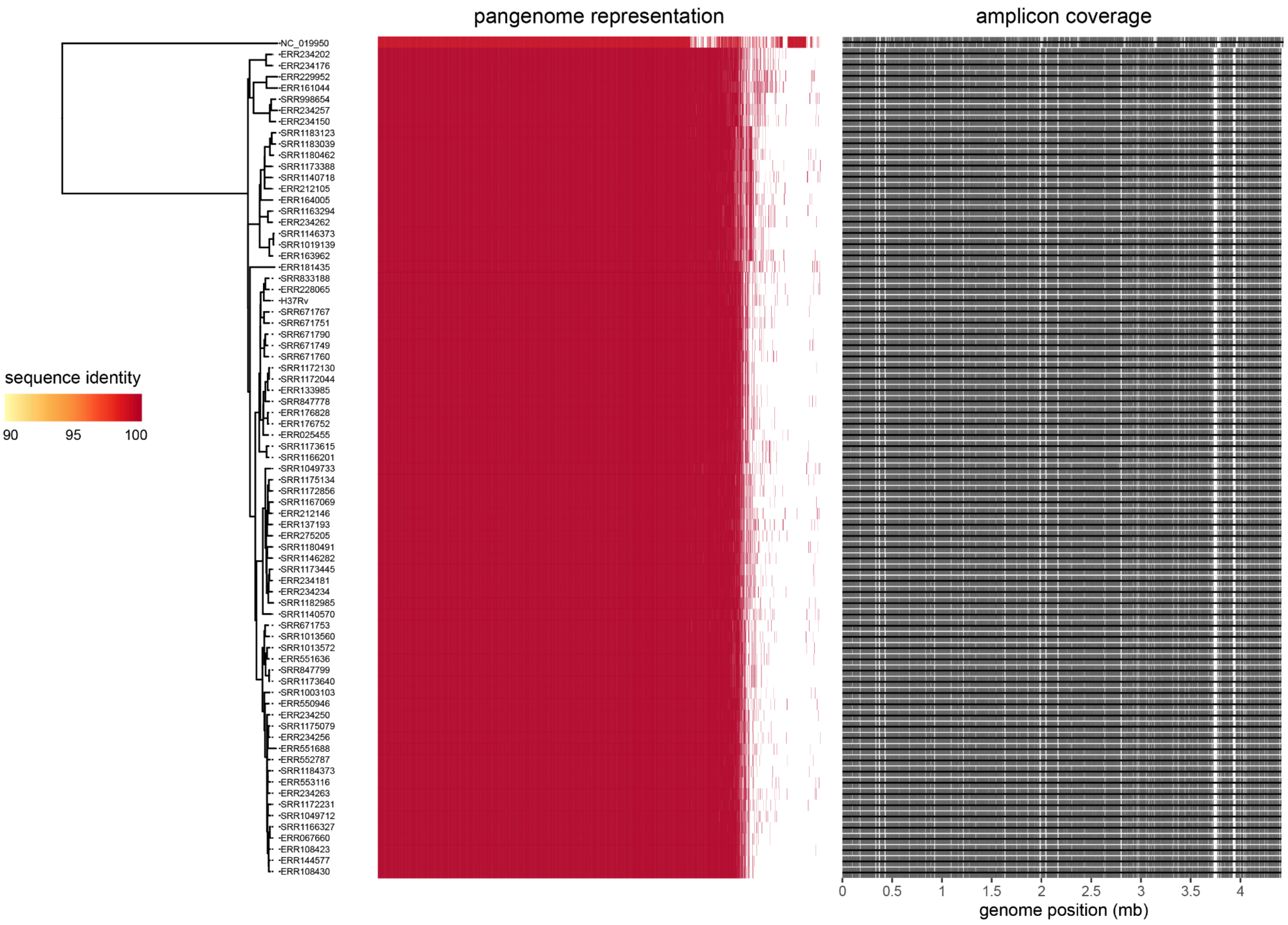
pangenome representation
amplicon coverage
sequence identity
90
95
100
NC_019950
ERR234202
ERR234176
ERR229952
ERR161044
SRR998654
ERR234257
ERR234150
SRR1183123
SRR1183039
SRR1180462
SRR1173388
SRR1140718
ERR212105
ERR164005
SRR1163294
ERR234262
SRR1146373
SRR1019139
ERR163962
ERR181435
SRR833188
ERR228065
H37Rv
SRR671767
SRR671751
SRR671790
SRR671749
SRR671760
SRR1172130
SRR1172044
ERR133985
SRR847778
ERR176828
ERR176752
ERR025455
SRR1173615
SRR1166201
SRR1049733
SRR1175134
SRR1172856
SRR1167069
ERR212146
ERR137193
ERR275205
SRR1180491
SRR1146282
SRR1173445
ERR234181
ERR234234
SRR1182985
SRR1140570
SRR671753
SRR1013560
SRR1013572
ERR551636
SRR847799
SRR1173640
SRR1003103
ERR550946
ERR234250
SRR1175079
ERR234256
ERR551688
ERR552787
SRR1184373
ERR553116
ERR234263
SRR1172231
SRR1049712
SRR1166327
ERR067660
ERR108423
ERR144577
ERR108430
0
0.5
1
1.5
2
2.5
3
3.5
4
genome position (mb)

**Figure S1A.** ***Streptococcus pneumoniae*** **pangenome.**

Pangenome representation of *S. pneumoniae* whole genome Global Pneumococcal Sequence Clusters (GPSC) sequences (n=35), *Streptococcus mitis* outgroup (Accession: AP023349), and reference *S. pneumoniae* used to build primers (Accession: NC_017592). Shaded bar graphs (middle) denote genes shared amongst clades, color denotes average nucleotide identity. Predicted amplicon coverage (right) is shown in grey with forward and reverse amplicon pairs displayed above and below the line.

**Figure S1B.** ***Mycobacterium tuberculosis*** **pangenome.**

Pangenome representation of *M. tuberculosis* whole genome sequences (n=76), *Mycobacterium canettii* outgroup (Accession: NC_019950), and reference *M. tuberculosis* sequence used to build primers (Accession: H37Rv). Shaded bar graphs (middle) denote genes shared amongst clades, color denotes average nucleotide identity. Predicted amplicon coverage (right) is shown in grey with forward and reverse amplicon pairs displayed above and below the line.

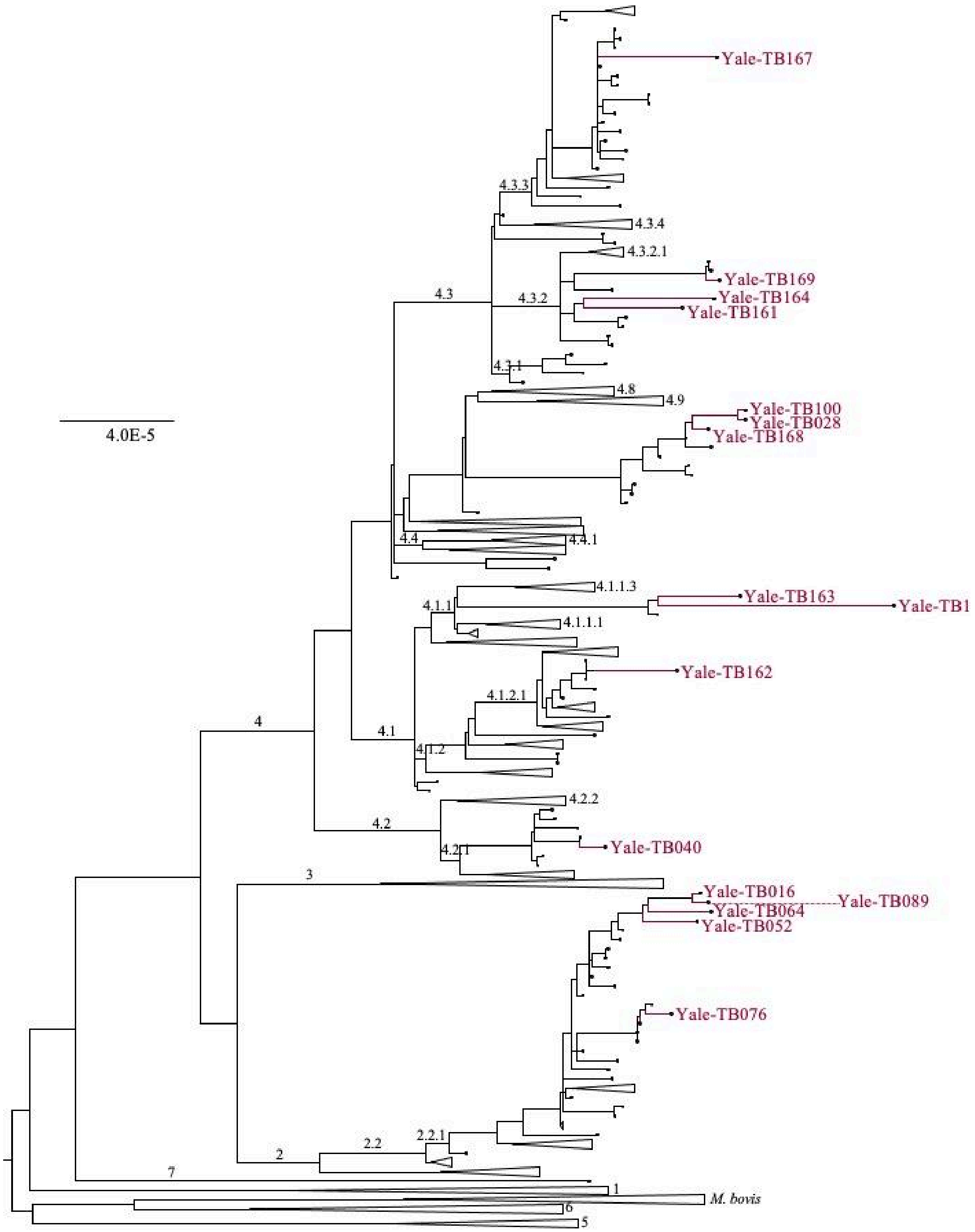


**Figure S2. Amplicon sequencing can be used in phylogenetic investigation.**
Maximum likelihood tree including all specimens with >80% coverage across the whole genome (highlighted in red), with sequences used for primer design (File S1A) as context. Branches are labeled according to lineage/sublineage according to a previously-described SNP barcoding system (Coll et al. 2014). For all samples, the primary lineage and sublineage called by Mykrobe is in accordance with the sublineage in the tree. A complete and interactive tree can be found at:
https://nextstrain.org/community/cck42/tbtrees@master/20241022

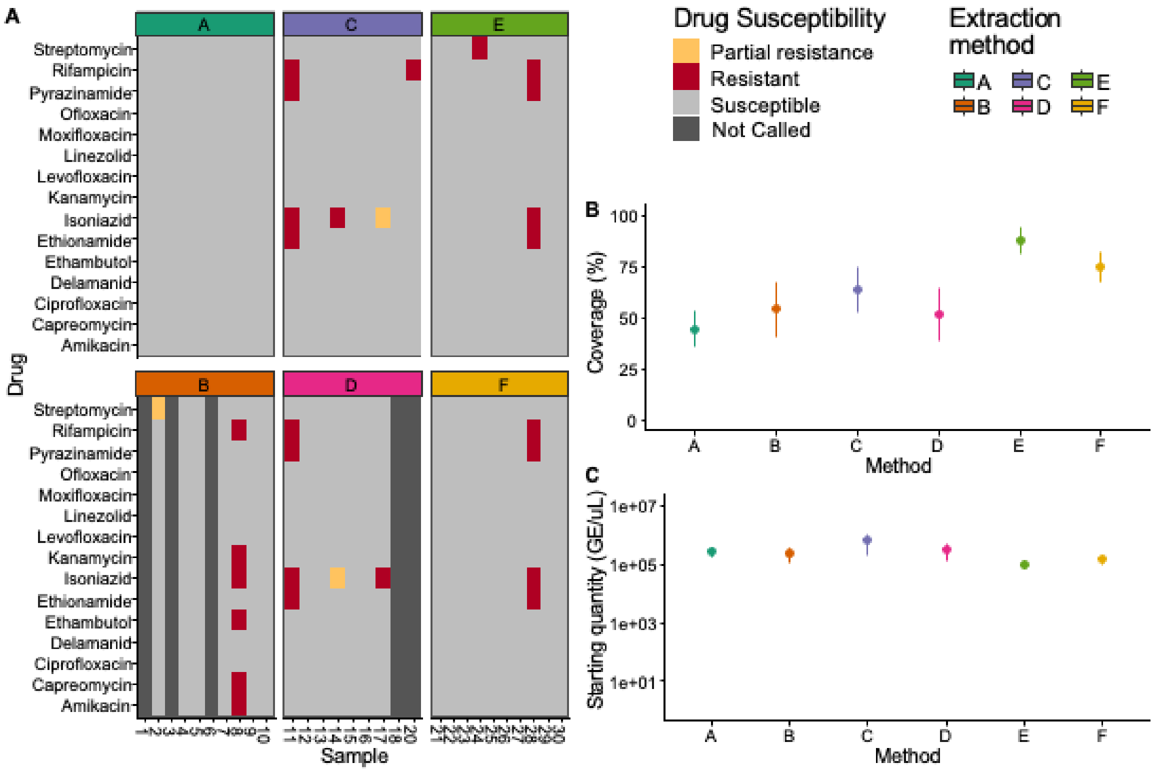


**Figure S3. Efficiency of extraction protocol affects antimicrobial resistance predictions.** Six different extraction protocols were assessed; 30 total unique specimens were extracted using 2 different protocols each. Mapping between patient and sample IDs is available in Table S1B. (A) Susceptibility to 14 anti-TB drugs by amplicon sequencing was predicted for all DNA extracted directly from sputum. (B) Comparison of mean coverage for each extraction protocol (dot) and standard error (line). (C) Comparison of mean starting quantity for each extraction protocol (dot) and standard error (line). Method E produced the highest average genome coverage and starting quantity with the least variation.

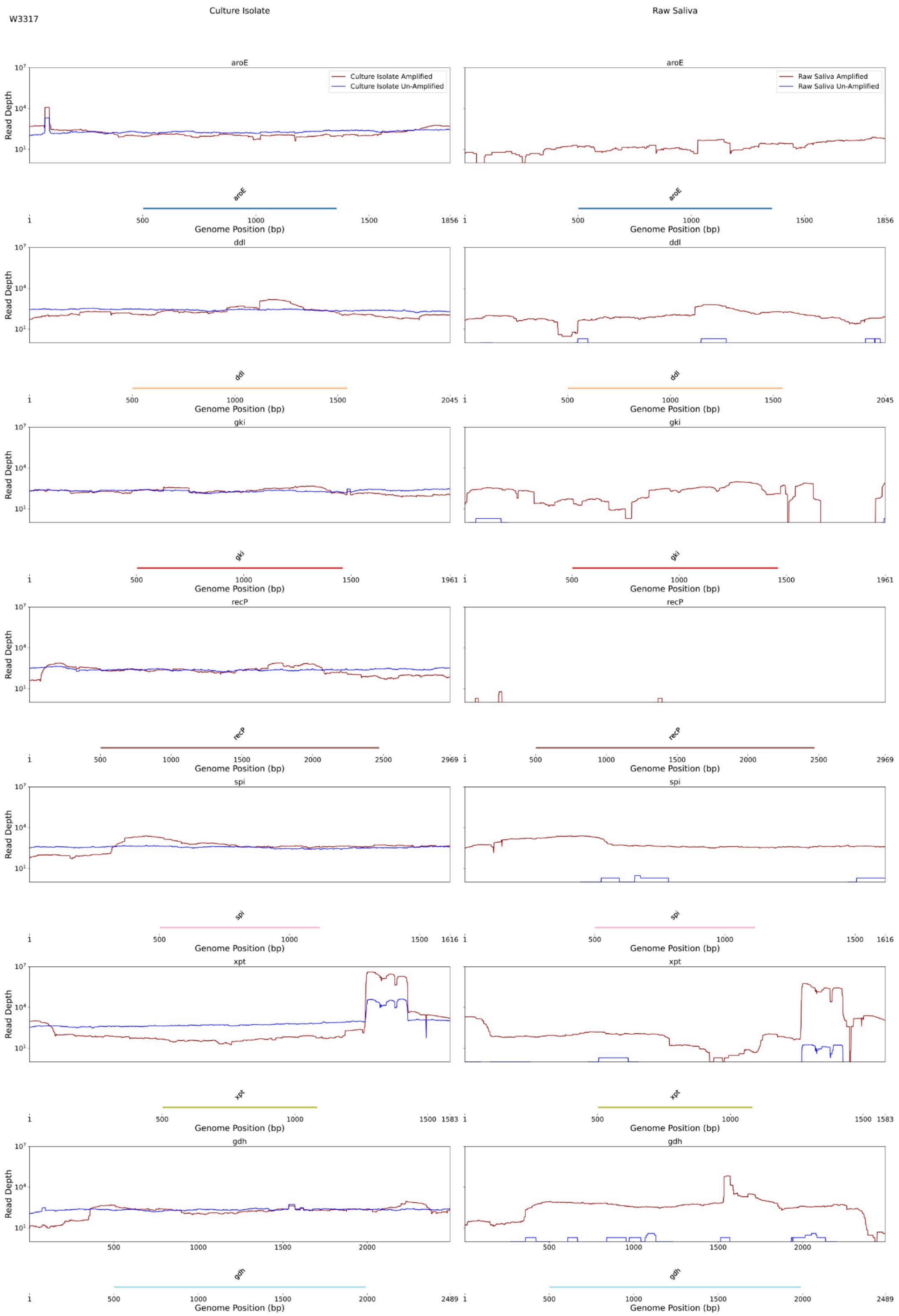
W3317
Culture Isolate
Raw Saliva
aroE
Culture Isolate Amplified
Culture Isolate Un-Amplified
Raw Saliva Amplified
Raw Saliva Un-Amplified
Read Depth
Genome Position (bp)
ddl
gki
recP
spi
xpt
gdh

**Figure S4. Raw read alignments to *Streptococcus pneumoniae* serotype 3 housekeeping genes under amplified and unamplified sequencing conditions.**
Read depth across the 7 *S. pneumoniae* serotype 3 housekeeping genes from sequence OXC141 is shown, including 500bp of flanking regions upstream and downstream of each allele, for paired samples with and without tiling amplification. Sample W3317 includes paired culture isolate and raw saliva samples. For more information on all paired samples, refer to Table S1A**.**

**Table S1A.** ***S. pneumoniae*** **clinical specimens**

| Sample | Origin* | Culture-enriched saliva** | Raw saliva | Cultured isolate |
|---|---|---|---|---|
| A889 | YNHH Biorepository | x | x | |
| B042 | YNHH Biorepository | x | x | |
| C677 | YNHH Biorepository | x | x | |
| W1526 | YNHH Biorepository | x | x | |
| W1527 | YNHH Biorepository | x | x | x |
| W1695 | YNHH Biorepository | x | x | |
| W3317 | YNHH Biorepository | x | x | x |
| W3371 | YNHH Biorepository | x | x | |
| W4034 | YNHH Biorepository | x | x | x |
| B013 | YNHH Biorepository | x | x | |
| B595 | YNHH Biorepository | x | x | |
| B015 | YNHH Biorepository | x | x | |
| B687 | YNHH Biorepository | x | x | |
| W1075 | YNHH Biorepository | x | x | |
| W2987 | YNHH Biorepository | x | x | |
| CDC | CDC | | | x |
| H54 | Hungary | | | x |
| H74 | Hungary | | | x |
| H122 | Hungary | | | x |
| H136 | Hungary | | | x |
| H141 | Hungary | | | x |

*YNHH Biorepository specimens were remnants from asymptomatic healthcare workers and patients at Yale-New Haven Hospital (Wyllie et al. 2020). Hungary specimens were remnants collected from healthy children ages 3-6 in daycare centers in Hungary (Tóthpál et al. 2015).
**Culture-enriched specimens consist of raw saliva specimens cultured on blood agar supplemented with gentamicin in order to enrich for pneumococcus, as described previously (Wyllie et al. 2023)

**Table S1B.** ***M. tuberculosis*** **clinical specimens.**

| Sample | Patient Number | Type | Origin* | Heat | Liquefaction | Decontamination | Homogenization |
|---|---|---|---|---|---|---|---|
| 111-10300-18 | - | Cultured sputum | Moldova | - | NALC 0.5% | NaOH | - |
| 444-3403-18 | - | Cultured sputum | Moldova | - | NALC 0.5% | NaOH | - |
| 111-5264-18 | - | Cultured sputum | Moldova | - | NALC 0.5% | NaOH | - |
| 111-10712-18 | - | Cultured sputum | Moldova | - | NALC 0.5% | NaOH | - |
| 111-2565-18 | - | Cultured sputum | Moldova | - | NALC 0.5% | NaOH | - |
| 444-2261-18 | - | Cultured sputum | Moldova | - | NALC 0.5% | NaOH | - |
| 444-2281-18 | - | Cultured sputum | Moldova | - | NALC 0.5% | NaOH | - |
| 444-2588-18 | - | Cultured sputum | Moldova | - | NALC 0.5% | NaOH | - |
| Peru-1 | 1 | Sputum | Peru | 95℃ x 20 min | Saponin | None | 5.5m/s x 40s x 2 cycles |
| Peru-2 | 2 | Sputum | Peru | 95℃ x 20 min | Saponin | None | 5.5m/s x 40s x 2 cycles |
| Peru-3 | 3 | Sputum | Peru | 95℃ x 20 min | Saponin | None | 5.5m/s x 40s x 2 cycles |
| Peru-4 | 4 | Sputum | Peru | 95℃ x 20 min | Saponin | None | 5.5m/s x 40s x 2 cycles |
| Peru-5 | 5 | Sputum | Peru | 95℃ x 20 min | Saponin | None | 5.5m/s x 40s x 2 cycles |
| Peru-6 | 6 | Sputum | Peru | 95℃ x 20 min | Saponin | None | 5.5m/s x 40s x 2 cycles |
| Peru-7 | 7 | Sputum | Peru | 95℃ x 20 min | Saponin | None | 5.5m/s x 40s x 2 cycles |
| Peru-8 | 8 | Sputum | Peru | 95℃ x 20 min | Saponin | None | 5.5m/s x 40s x 2 cycles |
| Peru-9 | 9 | Sputum | Peru | 95℃ x 20 min | Saponin | None | 5.5m/s x 40s x 2 cycles |
| Peru-10 | 10 | Sputum | Peru | 95℃ x 20 min | Saponin | None | 5.5m/s x 40s x 2 cycles |
| Peru-11 | 1 | Sputum | Peru | 95℃ x 20 min | NALC 0.5% | None | 5.5m/s x 40s x 2 cycles |
| Peru-12 | 2 | Sputum | Peru | 95℃ x 20 min | NALC 0.5% | None | 5.5m/s x 40s x 2 cycles |
| Peru-13 | 3 | Sputum | Peru | 95℃ x 20 min | NALC 0.5% | None | 5.5m/s x 40s x 2 cycles |

| | | | | | | | |
|---|---|---|---|---|---|---|---|
| Peru-14 | 4 | Sputum | Peru | 95℃ x 20 min | NALC 0.5% | None | 5.5m/s x 40s x 2 cycles |
| Peru-15 | 5 | Sputum | Peru | 95℃ x 20 min | NALC 0.5% | None | 5.5m/s x 40s x 2 cycles |
| Peru-16 | 6 | Sputum | Peru | 95℃ x 20 min | NALC 0.5% | None | 5.5m/s x 40s x 2 cycles |
| Peru-17 | 7 | Sputum | Peru | 95℃ x 20 min | NALC 0.5% | None | 5.5m/s x 40s x 2 cycles |
| Peru-18 | 8 | Sputum | Peru | 95℃ x 20 min | NALC 0.5% | None | 5.5m/s x 40s x 2 cycles |
| Peru-19 | 9 | Sputum | Peru | 95℃ x 20 min | NALC 0.5% | None | 5.5m/s x 40s x 2 cycles |
| Peru-20 | 10 | Sputum | Peru | 95℃ x 20 min | NALC 0.5% | None | 5.5m/s x 40s x 2 cycles |
| Peru-21 | 11 | Sputum | Peru | None | NALC 1% | None | 5.5m/s x 40s x 2 cycles |
| Peru-22 | 12 | Sputum | Peru | None | NALC 1% | None | 5.5m/s x 40s x 2 cycles |
| Peru-23 | 13 | Sputum | Peru | None | NALC 1% | None | 5.5m/s x 40s x 2 cycles |
| Peru-24 | 14 | Sputum | Peru | None | NALC 1% | None | 5.5m/s x 40s x 2 cycles |
| Peru-25 | 15 | Sputum | Peru | None | NALC 1% | None | 5.5m/s x 40s x 2 cycles |
| Peru-26 | 16 | Sputum | Peru | None | NALC 1% | None | 5.5m/s x 40s x 2 cycles |
| Peru-27 | 17 | Sputum | Peru | None | NALC 1% | None | 5.5m/s x 40s x 2 cycles |
| Peru-28 | 18 | Sputum | Peru | None | NALC 1% | None | 5.5m/s x 40s x 2 cycles |
| Peru-29 | 19 | Sputum | Peru | None | NALC 1% | None | 5.5m/s x 40s x 2 cycles |
| Peru-30 | 20 | Sputum | Peru | None | NALC 1% | None | 5.5m/s x 40s x 2 cycles |
| Peru-31 | 11 | Sputum | Peru | None | NALC 2% | None | 5.5m/s x 40s x 2 cycles |
| Peru-32 | 12 | Sputum | Peru | None | NALC 2% | None | 5.5m/s x 40s x 2 cycles |
| Peru-33 | 13 | Sputum | Peru | None | NALC 2% | None | 5.5m/s x 40s x 2 cycles |
| Peru-34 | 14 | Sputum | Peru | None | NALC 2% | None | 5.5m/s x 40s x 2 cycles |
| Peru-35 | 15 | Sputum | Peru | None | NALC 2% | None | 5.5m/s x 40s x 2 cycles |
| Peru-36 | 16 | Sputum | Peru | None | NALC 2% | None | 5.5m/s x 40s x 2 cycles |

| | | | | | | | |
|---|---|---|---|---|---|---|---|
| Peru-37 | 17 | Sputum | Peru | None | NALC 2% | None | 5.5m/s x 40s x 2 cycles |
| Peru-38 | 18 | Sputum | Peru | None | NALC 2% | None | 5.5m/s x 40s x 2 cycles |
| Peru-39 | 19 | Sputum | Peru | None | NALC 2% | None | 5.5m/s x 40s x 2 cycles |
| Peru-40 | 20 | Sputum | Peru | None | NALC 2% | None | 5.5m/s x 40s x 2 cycles |
| Peru-41 | 21 | Sputum | Peru | None | NALC 0.5% | NaOH | 5.5m/s x 40s x 2 cycles |
| Peru-42 | 22 | Sputum | Peru | None | NALC 0.5% | NaOH | 5.5m/s x 40s x 2 cycles |
| Peru-43 | 23 | Sputum | Peru | None | NALC 0.5% | NaOH | 5.5m/s x 40s x 2 cycles |
| Peru-44 | 24 | Sputum | Peru | None | NALC 0.5% | NaOH | 5.5m/s x 40s x 2 cycles |
| Peru-45 | 25 | Sputum | Peru | None | NALC 0.5% | NaOH | 5.5m/s x 40s x 2 cycles |
| Peru-46 | 26 | Sputum | Peru | None | NALC 0.5% | NaOH | 5.5m/s x 40s x 2 cycles |
| Peru-47 | 27 | Sputum | Peru | None | NALC 0.5% | NaOH | 5.5m/s x 40s x 2 cycles |
| Peru-48 | 28 | Sputum | Peru | None | NALC 0.5% | NaOH | 5.5m/s x 40s x 2 cycles |
| Peru-49 | 29 | Sputum | Peru | None | NALC 0.5% | NaOH | 5.5m/s x 40s x 2 cycles |
| Peru-50 | 30 | Sputum | Peru | None | NALC 0.5% | NaOH | 5.5m/s x 40s x 2 cycles |
| Peru-51 | 21 | Sputum | Peru | None | NALC 0.5% | NaOH | 5.5m/s x 20s x 2 cycles |
| Peru-52 | 22 | Sputum | Peru | None | NALC 0.5% | NaOH | 5.5m/s x 20s x 2 cycles |
| Peru-53 | 23 | Sputum | Peru | None | NALC 0.5% | NaOH | 5.5m/s x 20s x 2 cycles |
| Peru-54 | 24 | Sputum | Peru | None | NALC 0.5% | NaOH | 5.5m/s x 20s x 2 cycles |
| Peru-55 | 25 | Sputum | Peru | None | NALC 0.5% | NaOH | 5.5m/s x 20s x 2 cycles |
| Peru-56 | 26 | Sputum | Peru | None | NALC 0.5% | NaOH | 5.5m/s x 20s x 2 cycles |
| Peru-57 | 27 | Sputum | Peru | None | NALC 0.5% | NaOH | 5.5m/s x 20s x 2 cycles |
| Peru-58 | 28 | Sputum | Peru | None | NALC 0.5% | NaOH | 5.5m/s x 20s x 2 cycles |
| Peru-59 | 29 | Sputum | Peru | None | NALC 0.5% | NaOH | 5.5m/s x 20s x 2 cycles |

| | | | | | | | |
|---|---|---|---|---|---|---|---|
| Peru-60 | 30 | Sputum | Peru | None | NALC 0.5% | NaOH | 5.5m/s x 20s x 2 cycles |

*All samples from Moldova were remnant samples of DNA extracted from culture for a previous study (Yang et al. 2022). Peru specimens were extracted at Cayetano University, Peru.
**Cetyltrimethylammonium bromide (CTAB) method was used as described previously (Schiebelhut et al. 2017).

**Table S2. Samples used to predict off-target amplification.**

| Pathogen | Sample ID | Serotype List | Predicted Coverage |
|---|---|---|---|
| *S. mitis* | AP023349 | | 32.18% |
| *S. pneumoniae* | NC_017592 | | 98.93% |
| *S. pneumoniae* | GPSC3 | 8,33F,11A,22F,18C,3,15A,33A,23F,31 | 81.44% |
| *S. pneumoniae* | GPSC4 | 19A,15BC,14,19B,19F | 81.60% |
| *S. pneumoniae* | GPSC8 | 5 | 82.74% |
| *S. pneumoniae* | GPSC15 | 7F,19F | 88.50% |
| *S. pneumoniae* | GPSC21 | 19F,19A,14 | 82.07% |
| *S. pneumoniae* | GPSC22 | 11A,15A,20A,9V,35A,19A,19F,6A | 84.29% |
| *S. pneumoniae* | GPSC26 | 12F,46,12A,40,9V | 81.38% |
| *S. pneumoniae* | GPSC31 | 1 | 82.32% |
| *S. pneumoniae* | GPSC32 | 12F,7F,8,9N | 85.34% |
| *S. pneumoniae* | GPSC34 | 34,22F,23F,15A | 85.13% |
| *S. pneumoniae* | GPSC37 | 6B,23F | 82.08% |
| *S. pneumoniae* | GPSC40 | 15BC,22A,18C,10A,17F,15A,23F,11A,6B | 82.00% |
| *M. canettii* | NC_019950 | | 89.44% |
| *M. tuberculosis* | H37Rv | | 94.30% |
| *M. tuberculosis* | SRR1173640 | | 94.26% |
| *M. tuberculosis* | ERR144577 | | 94.27% |
| *M. tuberculosis* | ERR212146 | | 94.26% |
| *M. tuberculosis* | SRR1172044 | | 94.30% |
| *M. tuberculosis* | SRR671749 | | 94.30% |
| *M. tuberculosis* | ERR181435 | | 94.30% |
| *M. tuberculosis* | SRR1180462 | | 94.23% |
| *M. tuberculosis* | SRR1019139 | | 94.26% |
| *M. tuberculosis* | SRR1163294 | | 94.24% |
| *M. tuberculosis* | SRR998654 | | 94.24% |
| *M. tuberculosis* | ERR161044 | | 94.26% |
| *M. tuberculosis* | ERR234202 | | 94.25% |

**Table S3. PneumoKITy analysis of *S. pneumoniae.***

| Sequence ID | MASH Hit Kmer Percentage | Predicted Serotype | Sample Type / Paired ID |
|---|---|---|---|
| CS00030 | {'03': 95.2, '01': 17.9, '36': 17.5, '09L': 16.5, '09N': 16.1} | 3 | Culture Isolate - CDC |
| CS00032 | {'03': 44.4, '36': 17.8, '20': 16.8, '09L': 16.6, '09N': 16.3} | Below 70% hit - Poor Sequence quality, variant or non-typeable organisms. | Culture Isolate - H54 |
| CS00034 | {'03': 96.0, '01': 18.4, '36': 17.4, '09L': 16.9, '09N': 16.5} | 3 | Culture Isolate - H74 |
| CS00036 | {'03': 95.6, '01': 18.3, '36': 17.2, '09L': 16.2, '20': 15.6} | 3 | Culture Isolate - H122 |
| CS00038 | {'03': 94.8, '01': 17.3, '36': 16.8, '09L': 15.6, '09N': 15.3} | 3 | Culture Isolate - H136 |
| CS00040 | {'03': 95.0, '36': 21.5, '01': 20.4, '20': 20.0, '09L': 18.3} | 3 | Culture Isolate - H141 |
| CS00153 | {'03': 16.1, '01': 5.8, '45': 5.5, '09L': 5.1, '22F': 5.0} | Below 20% hit - inadequate DNA or acapsular organism, check species identity and sequence quality. | Culture Enriched - A889 |
| CS00154 | {'09L': 8.4, '09N': 8.0, '03': 6.5, '01': 5.8, '22F': 5.4} | Below 20% hit - inadequate DNA or acapsular organism, check species identity and sequence quality. | Culture Enriched - B042 |
| CS00155 | {'03': 6.0, '01': 5.0, '45': 4.5, '25F': 3.7, '35B': 3.7} | Below 20% hit - inadequate DNA or acapsular organism, check species identity and sequence quality. | Culture Enriched - C677 |
| CS00156 | {'01': 5.2, '22F': 4.5, '09L': 4.0, '09N': 3.8, '38': 3.7} | Below 20% hit - inadequate DNA or acapsular organism, check species identity and sequence quality. | Culture Enriched - W1526 |
| CS00157 | {'01': 6.0, '45': 3.0, '09N': 2.7, '25F': 2.5, '09L': 2.4} | Below 20% hit - inadequate DNA or acapsular organism, check species identity and sequence quality. | Culture Enriched - W1527 |
| CS00158 | {'01': 6.6, '22F': 6.0, '45': 5.1, '18A': 4.3, '25F': 4.2} | Below 20% hit - inadequate DNA or acapsular organism, check species identity and sequence quality. | Culture Enriched - W1695 |
| CS00159 | {'03': 12.4, '45': 6.0, '01': 5.1, '35B': 5.0, '18A': 4.9} | Below 20% hit - inadequate DNA or acapsular organism, check species identity and sequence quality. | Culture Enriched - W3317 |

| | | | |
|---|---|---|---|
| CS00160 | {'01': 5.3, '22F': 4.5, '45': 4.4, '25F': 4.3, '25A': 4.2} | Below 20% hit - inadequate DNA or acapsular organism, check species identity and sequence quality. | Culture Enriched - W3371 |
| CS00161 | {'01': 6.0, '22F': 4.9, '25F': 4.6, '35B': 4.5, '09L': 4.4} | Below 20% hit - inadequate DNA or acapsular organism, check species identity and sequence quality. | Culture Enriched - W4034 |
| CS00162 | {'03': 56.7, '01': 13.4, '09L': 12.4, '09N': 12.2, '36': 11.5} | Below 70% hit - Poor Sequence quality, variant or non-typeable organisms. | Saliva - A889 |
| CS00163 | {'01': 5.7, '35B': 4.7, '09L': 4.3, '38': 4.3, '25F': 4.2} | Below 20% hit - inadequate DNA or acapsular organism, check species identity and sequence quality. | Saliva - B042 |
| CS00164 | {'03': 39.9, '01': 11.0, '09L': 9.4, '09N': 9.2, '18A': 8.7} | Below 70% hit - Poor Sequence quality, variant or non-typeable organisms. | Saliva - C677 |
| CS00165 | {'03': 5.8, '01': 5.0, '25F': 4.1, '38': 4.1, '25A': 4.0} | Below 20% hit - inadequate DNA or acapsular organism, check species identity and sequence quality. | Saliva - W1526 |
| CS00166 | {'01': 8.5, '09L': 5.8, '09N': 5.1, '14': 4.2, '22F': 4.1} | Below 20% hit - inadequate DNA or acapsular organism, check species identity and sequence quality. | Saliva - W1527 |
| CS00167 | {'03': 8.5, '45': 6.1, '01': 5.5, '09L': 5.0, '22F': 5.0} | Below 20% hit - inadequate DNA or acapsular organism, check species identity and sequence quality. | Saliva - W1695 |
| CS00168 | {'01': 10.2, '09L': 8.2, '09N': 7.4, '45': 6.3, '18A': 6.0} | Below 20% hit - inadequate DNA or acapsular organism, check species identity and sequence quality. | Saliva - W3317 |
| CS00169 | {'01': 8.1, '45': 6.1, '09L': 4.6, '22F': 4.3, '09N': 4.2} | Below 20% hit - inadequate DNA or acapsular organism, check species identity and sequence quality. | Saliva - W3371 |
| CS00170 | {'03': 13.7, '01': 6.7, '45': 6.5, '09L': 5.2, '09N': 4.8} | Below 20% hit - inadequate DNA or acapsular organism, check species identity and sequence quality. | Saliva - W4034 |
| CS00174 | {'03': 95.6, '01': 18.5, '36': 17.9, '09L': 16.9, '20': 16.6} | 3 | Culture Isolate - W1527 |

| | | | |
|---|---|---|---|
| CS00175 | {'03': 94.8, '01': 18.4, '36': 17.9, '09L': 17.0, '20': 16.6} | 3 | Culture Isolate - W3317 |
| CS00176 | {'03': 95.6, '01': 18.1, '36': 18.1, '09L': 16.9, '09N': 16.6} | 3 | Culture Isolate - W4034 |
| CS00177 | {'01': 3.9, '09L': 2.2, '25F': 1.6, '35B': 1.6, '38': 1.6} | Below 20% hit - inadequate DNA or acapsular organism, check species identity and sequence quality. | Saliva - B013 |
| CS00178 | {'01': 6.9, '09L': 3.9, '20': 3.6, '09N': 3.3, '22F': 3.1} | Below 20% hit - inadequate DNA or acapsular organism, check species identity and sequence quality. | Saliva - B595 |
| CS00179 | {'35B': 5.9, '06A-I': 5.4, '34': 5.3, '06A-III': 5.2, '06B-I': 5.2} | Below 20% hit - inadequate DNA or acapsular organism, check species identity and sequence quality. | Saliva - B015 |
| CS00180 | {'01': 5.1, '09N': 1.8, '25A': 1.8, '25F': 1.8, '38': 1.8} | Below 20% hit - inadequate DNA or acapsular organism, check species identity and sequence quality. | Saliva - W2987 |
| CS00181 | {'01': 2.2, '06A-I': 1.3, '22F': 0.9, '17F': 0.8, '34': 0.8} | Below 20% hit - inadequate DNA or acapsular organism, check species identity and sequence quality. | Saliva - B687 |
| CS00182 | {'01': 5.8, '25F': 3.8, '25A': 3.7, '09L': 3.6, '22F': 3.4} | Below 20% hit - inadequate DNA or acapsular organism, check species identity and sequence quality. | Saliva - W1075 |
| CS00186 | {'01': 3.2, '24F': 2.7, '24B': 2.6, '24F2': 2.5, '18A': 2.4} | Below 20% hit - inadequate DNA or acapsular organism, check species identity and sequence quality. | Culture Enriched - B595 |
| CS00187 | {'01': 4.8, '22F': 2.1, '38': 1.4, '10A': 1.3, '25A': 1.3} | Below 20% hit - inadequate DNA or acapsular organism, check species identity and sequence quality. | Culture Enriched - B013 |
| CS00188 | {'01': 4.5, '14': 2.0, '22F': 2.0, '09L': 1.9, '20': 1.9} | Below 20% hit - inadequate DNA or acapsular organism, check species identity and sequence quality. | Culture Enriched - B015 |
| CS00189 | {'01': 5.4, '22F': 4.8, '25F': 4.6, '25A': 4.4, '38': 4.3} | Below 20% hit - inadequate DNA or acapsular organism, check species identity and sequence quality. | Culture Enriched - W1075 |

| | | | |
|---|---|---|---|
| CS00190 | {'01': 4.9, '25F': 3.4, '25A': 3.3, '38': 3.3, '09N': 3.1} | Below 20% hit - inadequate DNA or acapsular organism, check species identity and sequence quality. | Culture Enriched - W2987 |
| CS00191 | {'01': 3.3, '06A-I': 1.1, '03': 0.9, '22F': 0.8, '20': 0.7} | Below 20% hit - inadequate DNA or acapsular organism, check species identity and sequence quality. | Culture Enriched - B687 |

Output from PneumoKITy (Pneumococcal Kmer Integrated Typing) serotyping tool. The predicted phenotypic serotype reflects only first-stage determinations by PnuemoKITy. MASH k-mer hit percentages for each sample's top five serotype hits are listed from highest to lowest. For more information on all paired samples, refer to Table S1A**.**

**Table S4. MLST analysis of *S. pneumoniae* culture isolates.**

| Sequence ID | PubMLST Scheme Name | Sequence Type | aroE | gdh | gki | recP | spi | xpt | ddl |
|---|---|---|---|---|---|---|---|---|---|
| CS00030 | *S. pneumoniae* | 180 | 7 | 15 | 2 | 10 | 6 | 1 | 22 |
| CS00032 | *S. pneumoniae* | | 2 | 8 | 263? | 4 | 6 | 1075? | 1 |
| CS00034 | *S. pneumoniae* | 180 | 7 | 15 | 2 | 10 | 6 | 1 | 22 |
| CS00036 | *S. pneumoniae* | 180 | 7 | 15 | 2 | 10 | 6 | 1 | 22 |
| CS00038 | *S. pneumoniae* | 180 | 7 | 15 | 2 | 10 | 6 | 1 | 22 |
| CS00040 | *S. pneumoniae* | 180 | 7 | 15 | 2 | 10 | 6 | 1 | 22 |
| CS00174 | *S. pneumoniae* | | 7 | 15 | 2 | 10 | 6 | 1 | ~893 |
| CS00175 | | | | | | | | | |
| CS00176 | *S. pneumoniae* | | 602? | 15 | 2 | 10 | 6 | - | 22 |

Output from PubMLST typing scheme search using amplicon data. Exact allele matches are listed as whole numbers, partial allele matches are denoted with a question mark (e.g. n?), novel full-length alleles are denoted with a tilde (e.g. ~n), and missing alleles are denoted with a dash (e.g. -). Undetermined information is left blank. MLST typing schemes could not be determined for raw saliva and culture-enriched saliva.

**Table S5. Antibiotic resistance in *S. pneumoniae* samples.**

| Sequence ID | START | END | GENE | % COV | % ID | ACCESSION | RESISTANCE | Sample ID | Sample Type |
|---|---|---|---|---|---|---|---|---|---|
| CS00030 | 4546 | 6312 | patB | 100 | 99.77 | AE005672.3:1982324-1980557 | fluoroquinolone | CDC | Culture Isolate |
| CS00030 | 7087 | 8781 | patA | 100 | 99.65 | AE005672.3:1984810-1983115 | fluoroquinolone | CDC | Culture Isolate |
| CS00030 | 14 | 862 | RlmA(II) | 100 | 99.53 | CP007593.1:2148923-2149772 | lincosamide;macrolide | CDC | Culture Isolate |
| CS00032 | 7901 | 9100 | pmrA | 100 | 99.67 | AE007317.1:866210-867410 | fluoroquinolone | H54 | Culture Isolate |
| CS00032 | 4694 | 6460 | patB | 100 | 99.6 | AE005672.3:1982324-1980557 | fluoroquinolone | H54 | Culture Isolate |
| CS00032 | 128 | 976 | RlmA(II) | 100 | 100 | CP007593.1:2148923-2149772 | lincosamide;macrolide | H54 | Culture Isolate |
| CS00032 | 3125 | 4819 | patA | 100 | 99.41 | AE005672.3:1984810-1983115 | fluoroquinolone | H54 | Culture Isolate |
| CS00034 | 19872 | 20720 | RlmA(II) | 100 | 99.53 | CP007593.1:2148923-2149772 | lincosamide;macrolide | H74 | Culture Isolate |
| CS00034 | 4546 | 6312 | patB | 100 | 99.77 | AE005672.3:1982324-1980557 | fluoroquinolone | H74 | Culture Isolate |
| CS00034 | 7087 | 8781 | patA | 100 | 99.65 | AE005672.3:1984810-1983115 | fluoroquinolone | H74 | Culture Isolate |
| CS00036 | 4546 | 6312 | patB | 100 | 99.77 | AE005672.3:1982324-1980557 | fluoroquinolone | H122 | Culture Isolate |
| CS00036 | 35 | 883 | RlmA(II) | 100 | 99.53 | CP007593.1:2148923-2149772 | lincosamide;macrolide | H122 | Culture Isolate |
| CS00036 | 4280 | 5974 | patA | 100 | 99.65 | AE005672.3:1984810-1983115 | fluoroquinolone | H122 | Culture Isolate |
| CS00038 | 14 | 862 | RlmA(II) | 100 | 99.53 | CP007593.1:2148923-2149772 | lincosamide;macrolide | H136 | Culture Isolate |
| CS00038 | 4546 | 6312 | patB | 100 | 99.77 | AE005672.3:1982324-1980557 | fluoroquinolone | H136 | Culture Isolate |
| CS00038 | 77 | 1771 | patA | 100 | 99.65 | AE005672.3:1984810-1983115 | fluoroquinolone | H136 | Culture Isolate |
| CS00040 | 5017 | 6216 | pmrA | 100 | 99.25 | AE007317.1:866210-867410 | fluoroquinolone | H141 | Culture Isolate |
| CS00040 | 20887 | 21735 | RlmA(II) | 100 | 99.53 | CP007593.1:2148923-2149772 | lincosamide;macrolide | H141 | Culture Isolate |
| CS00040 | 779 | 2473 | patA | 100 | 99.65 | AE005672.3:1984810-1983115 | fluoroquinolone | H141 | Culture Isolate |
| CS00040 | 4402 | 6168 | patB | 100 | 99.77 | AE005672.3:1982324-1980557 | fluoroquinolone | H141 | Culture Isolate |
| CS00157 | 63 | 807 | RlmA(II) | 87.75 | 85.77 | CP007593.1:2148923-2149772 | lincosamide;macrolide | W1527 | Culture Enriched Saliva |

| | | | | | | | | | |
|---|---|---|---|---|---|---|---|---|---|
| CS00166 | 1 | 1391 | patB | 78.72 | 94.46 | AE005672.3:1982324-1980557 | fluoroquinolone | W1527 | Raw Saliva |
| CS00166 | 148 | 988 | RlmA(II) | 98.94 | 84.32 | CP007593.1:2148923-2149772 | lincosamide;macrolide | W1527 | Raw Saliva |
| CS00166 | 186 | 884 | RlmA(II) | 82.33 | 90.42 | CP007593.1:2148923-2149772 | lincosamide;macrolide | W1527 | Raw Saliva |
| CS00170 | 3186 | 4880 | patA | 100 | 94.22 | AE005672.3:1984810-1983115 | fluoroquinolone | W4034 | Raw Saliva |
| CS00170 | 4894 | 6660 | patB | 100 | 93.78 | AE005672.3:1982324-1980557 | fluoroquinolone | W4034 | Raw Saliva |
| CS00170 | 148 | 996 | RlmA(II) | 100 | 87.75 | CP007593.1:2148923-2149772 | lincosamide;macrolide | W4034 | Raw Saliva |
| CS00170 | 5281 | 6129 | RlmA(II) | 100 | 84.33 | CP007593.1:2148923-2149772 | lincosamide;macrolide | W4034 | Raw Saliva |
| CS00174 | 2259 | 3107 | RlmA(II) | 100 | 99.41 | CP007593.1:2148923-2149772 | lincosamide;macrolide | W1527 | Culture Isolate |
| CS00174 | 7512 | 8571 | pmrA | 88.33 | 99.25 | AE007317.1:866210-867410 | fluoroquinolone | W1527 | Culture Isolate |
| CS00174 | 1174 | 2940 | patB | 100 | 99.77 | AE005672.3:1982324-1980557 | fluoroquinolone | W1527 | Culture Isolate |
| CS00174 | 3715 | 5409 | patA | 100 | 99.65 | AE005672.3:1984810-1983115 | fluoroquinolone | W1527 | Culture Isolate |
| CS00176 | 1587 | 2435 | RlmA(II) | 100 | 99.53 | CP007593.1:2148923-2149772 | lincosamide;macrolide | W4034 | Culture Isolate |
| CS00176 | 836 | 2602 | patB | 100 | 99.77 | AE005672.3:1982324-1980557 | fluoroquinolone | W4034 | Culture Isolate |
| CS00176 | 3377 | 5071 | patA | 100 | 99.65 | AE005672.3:1984810-1983115 | fluoroquinolone | W4034 | Culture Isolate |

Output from ABRicate screening of contigs for antimicrobial resistance from various *S. pneumoniae* sample types.

**Table S6.** ***S. pneumoniae*** **qPCR results**

| Sample | Sample Type | lytA | piaB | Single Plex | Sample Type | lytA | piaB | Single Plex | Sample Type | lytA | piaB | Single Plex |
|---|---|---|---|---|---|---|---|---|---|---|---|---|
| A889 | Culture-Enriched Saliva | 26.13 | 26.58 | 25.22 | Raw Saliva | 27.05 | 27.68 | 25.97 | | | | |
| B042 | Culture-Enriched Saliva | 27.95 | 28.49 | 27.16 | Raw Saliva | 32.2 | 32.82 | 30.86 | | | | |
| C677 | Culture-Enriched Saliva | 25.13 | 26.12 | 24.67 | Raw Saliva | 29.9 | 29.84 | 28.78 | | | | |
| W1526 | Culture-Enriched Saliva | 30.92 | 32.3 | 30.63 | Raw Saliva | 36.72 | 38.27 | 37.56 | | | | |
| W1527 | Culture-Enriched Saliva | NaN | 35.38 | 34.72 | Raw Saliva | NaN | 37.81 | 34.4 | Culture Isolate | 14.84 | 14.51 | 13.21 |
| W1695 | Culture-Enriched Saliva | 28.72 | 29.8 | 28.29 | Raw Saliva | 39.37 | 39.54 | 38.32 | | | | |
| W3317 | Culture-Enriched Saliva | NaN | 29.75 | 29.21 | Raw Saliva | 35.48 | 33.47 | 32.22 | Culture Isolate | 14.53 | 14.31 | 13.06 |
| W3371 | Culture-Enriched Saliva | 32.15 | 33.5 | 31.76 | Raw Saliva | 37.33 | 38.52 | 35.53 | | | | |
| W4034 | Culture-Enriched Saliva | 27.06 | 28.06 | 26.69 | Raw Saliva | 35.11 | 35.47 | 33.04 | Culture Isolate | 16.82 | 17.4 | 15.52 |

**Appendix S1.** ***M. tuberculosis*** **extraction methods from sputum**

Method A

In a 15-mL falcon tube, 1 mL of sputum was heated on a thermal block at 95℃ for 30 minutes to inactivate. Liquefaction was performed by adding 1 mL of saponin, followed by vortexing for 20 seconds and inverting the tube 4-5 times, then allowing the sample to stand at room temperature for a minimum of 15 minutes but no longer than 20 minutes. Next, 13 mL phosphate buffer (pH 6.8) was added and the tube vortexed and inverted to neutralize the sample and terminate decontamination/liquefaction. The sample was then centrifuged at 3000 g for 15 minutes and supernatant removed. 300 µL PBS was added to the remaining pellet and vortexed. 300 µL of the resuspended pellet was homogenized in 2 cycles of 5.5 m/s for 40 seconds each, placing the sample on ice for 5 minutes after each cycle. The homogenized sample was then centrifuged at 16,000 g for 10 minutes. The supernatant was then transferred to a new 1.5 mL tube and a 1:1 magnetic bead cleanup performed.

Method B

In a 15-mL falcon tube, 1 mL of sputum was heated on a thermal block at 95℃ for 30 minutes to inactivate. Liquefaction was performed by adding 1 mL of 0.5% NALC, followed by vortexing for 20 seconds and inverting the tube 4-5 times, then allowing the sample to stand at room temperature for a minimum of 15 minutes but no longer than 20 minutes. Next, 13 mL phosphate buffer (pH 6.8) was added and the tube vortexed and inverted to neutralize the sample and terminate decontamination/liquefaction. The sample was then centrifuged at 3000 g for 15 minutes and supernatant removed. 300 µL PBS was added to the remaining pellet and vortexed. 300 µL of the resuspended pellet was homogenized in 2 cycles of 5.5 m/s for 40 seconds each, placing the sample on ice for 5 minutes after each cycle. The homogenized sample was then centrifuged at 16,000 g for 10 minutes. The supernatant was then transferred to a new 1.5 mL tube and a 1:1 magnetic bead cleanup performed.

Method C

1 mL of sputum was added to a 15-mL falcon tube, then combined decontamination/liquefaction was performed by adding 1% NALC, followed by vortexing for 20 seconds and inverting the tube 4-5 times, then allowing the sample to stand at room temperature for a minimum of 15 minutes but no longer than 20 minutes. Next, a 13 mL phosphate buffer (pH 6.8) was added and the tube vortexed and inverted to neutralize the sample and terminate decontamination/liquefaction. The sample was then centrifuged at 3000 g for 15 minutes and supernatant removed. 300 µL PBS was added to the remaining pellet and vortexed. 300 µL of the resuspended pellet was homogenized in 2 cycles of 5.5 m/s for 40 seconds each, placing the sample on ice for 5 minutes after each cycle. The homogenized sample was then centrifuged at 16,000 g for 10 minutes. The supernatant was then transferred to a new 1.5 mL tube and a 1:1 magnetic bead cleanup performed.

Method D

1 mL of sputum was added to a 15-mL falcon tube, then combined decontamination/liquefaction was performed by adding 2% NALC, followed by vortexing for 20 seconds and inverting the tube 4-5 times, then allowing the sample to stand at room temperature for a minimum of 15 minutes but no longer than 20 minutes. Next, a 13 mL phosphate buffer (pH 6.8) was added and the tube vortexed and inverted to neutralize the sample and terminate decontamination/liquefaction. The sample was then centrifuged at

3000 g for 15 minutes and supernatant removed. 300 µL PBS was added to the remaining pellet and vortexed. 300 µL of the resuspended pellet was homogenized in 2 cycles of 5.5 m/s for 40 seconds each, placing the sample on ice for 5 minutes after each cycle. The homogenized sample was then centrifuged at 16,000 g for 10 minutes. The supernatant was then transferred to a new 1.5 mL tube and a 1:1 magnetic bead cleanup performed.

Method E

1 mL of sputum was added to a 15-mL falcon tube, then combined decontamination/liquefaction was performed by adding a solution of 4% NaOH/2.9% Na citrate/0.5% NALC followed by vortexing for 20 seconds and inverting the tube 4-5 times, then allowing the sample to stand at room temperature for a minimum of 15 minutes but no longer than 20 minutes. Next, a 13 mL phosphate buffer (pH 6.8) was added and the tube vortexed and inverted to neutralize the sample and terminate decontamination/liquefaction. The sample was then centrifuged at 3000 g for 15 minutes and supernatant removed. 300 µL PBS was added to the remaining pellet and vortexed. 300 µL of the resuspended pellet was homogenized in 2 cycles of 5.5 m/s for 40 seconds each, placing the sample on ice for 5 minutes after each cycle. The homogenized sample was then centrifuged at 16,000 g for 10 minutes. The supernatant was then transferred to a new 1.5 mL tube and a 1:1 magnetic bead cleanup performed.

Method F

1 mL of sputum was added to a 15-mL falcon tube, then combined decontamination/liquefaction was performed by adding a solution of 4% NaOH/2.9% Na citrate/0.5% NALC. followed by vortexing for 20 seconds and inverting the tube 4-5 times, then allowing the sample to stand at room temperature for a minimum of 15 minutes but no longer than 20 minutes. Next, a 13 mL phosphate buffer (pH 6.8) was added and the tube vortexed and inverted to neutralize the sample and terminate decontamination/liquefaction. The sample was then centrifuged at 3000 g for 15 minutes and supernatant removed. 300 µL PBS was added to the remaining pellet and vortexed. 300 µL of the resuspended pellet was homogenized in 2 cycles of 5.5 m/s for 20 seconds each, placing the sample on ice for 5 minutes after each cycle. The homogenized sample was then centrifuged at 16,000 g for 10 minutes. The supernatant was then transferred to a new 1.5 mL tube and a 1:1 magnetic bead cleanup performed.
